# Competing forms of protein-protein association and DNA binding exhibited by BrxC from the BREX phage restriction system

**DOI:** 10.64898/2026.04.09.717308

**Authors:** Alexander J. Kaiser, Jennifer J. Readshaw, Lindsey A. Doyle, Maria Puiu, Abigail Kelly, Sydney F. McGuire, Julieta Peralta Acosta, Duc Vu, Andrew Nelson, Darren L. Smith, Lidia Araújo-Bazán, Ernesto Arias-Palomo, Yvette A. Luyten, Barry L. Stoddard, Tim R. Blower, Brett K. Kaiser

## Abstract

Bacteriophage exclusion (BREX) defense systems restrict phage infection via inhibition of phage DNA replication, while also modifying and protecting the bacterial genome. Type I BREX systems encode six conserved proteins, including a site-specific DNA methyltransferase. Host methylation requires a subset of BREX proteins, whereas phage restriction generally requires them all, suggesting that distinct but overlapping complexes mediate these activities. Full details of the mechanism and regulation of BREX remains to be understood. Here, we characterize the behavior and structures of the conserved BrxC AAA+ ATPase protein. BrxC forms multiple competing assemblages – various self-associating multimers, as well as a complex with BrxB-PglZ – that can be uncoupled via distinct point mutations, leading to differing effects on host methylation versus phage restriction. BrxC’s self-association, as well as its ability to bind DNA, is regulated by ATP binding and hydrolysis; BrxA and BrxB appear to also regulate those behaviors. These collective results suggest that BrxC may play a key role in controlling the two activities of BREX, with BrxB, BrxC and PglZ forming a core complex, and the equilibrium among competing assemblies containing those proteins modulating the balance between idling and activated restrictive states.

## INTRODUCTION

Anti-viral defense in bacteria, commonly termed ‘phage restriction’, was first described in the early 1950s, when two groups of investigators characterized the ability of different strains of a given bacterial species to resist replication of phage during laboratory infection experiments (1,2). The subsequent identification of mechanisms, pathways and factors that contribute to bacterial phage defense led to the identification of restriction endonucleases and their corresponding methyltransferase enzymes (i.e. ‘restriction-modification’ or ‘RM’ systems) (3–5), followed by the identification of CRISPR-based systems (6–10). Both operate via site-specific cleavage of DNA targets within the invading phage genome, and thereby serve as innate and adaptive defense systems, respectively.

Since the discovery of restriction endonucleases and CRISPR (and their subsequent development and application throughout biotechnology and biomedicine), many additional antiviral defense mechanisms and systems encoded in bacteria and archaea have been identified and characterized, as summarized in (11). Well over 100 such systems have been described; many were initially identified based on the location of their coding sequences within well-known genetic ‘defense islands’ associated with mobile genetic elements (12). These systems are notable for their diversity of form and function, ranging from those encoded by single genes and their corresponding products, to highly complex multi-gene operons, collectively establishing multiple layers of anti-viral defense (13). Their diverse mechanisms of action extend beyond degrading phage DNA, and include detection of phage-specific proteins and effectors, synthesis of bacterial second messengers and antiviral factors, inhibition of phage genome replication, and/or triggering of bacterial cell stasis leading to cell death (11,14).

One particularly complex group of phage restriction systems, found in roughly 10% of bacteria and archaea, is termed ‘Bacteriophage Exclusion’ or ‘BREX’ (15). BREX systems operate by preventing phage DNA replication within the infected bacterial host cell, while also relying on the action of a site-specific methyltransferase to protect the bacterial host genome from its action. Originally discovered in the 1980s and first named ‘Pgl’ based on its ‘**P**hage **G**rowth **L**imitation’ phenotype (16), BREX systems — currently classified into six types — comprise multiple genes (typically between 4 to 8) encoded within a single operon (15). All BREX systems are built around: (1) a sequence-specific DNA modifying enzyme (usually, but not always, a methyltransferase termed PglX or BrxX), and (2) two additional, strictly conserved factors: BrxC and PglZ (the latter also termed BrxZ). Recent studies have indicated that BrxA-and BrxB-like homologs are universally present across BREX subtypes, in some cases embedded within the polypeptides of other BREX proteins (17).

‘Type I’ BREX systems rely on the action of up to seven distinct proteins (**Figure 1a**), six of which have been individually characterized through structural, biochemical and functional approaches. Those factors include (i) a sequence-specific transcriptional regulator termed BrxR — present in a subset of BREX systems and widely associated with diverse defense systems — that appears to regulate the activity of the system and maintain it in a repressed ‘stand-by’ state prior to phage challenge (18–20); (ii) a sequence-specific DNA methyltransferase, termed PglX or BrxX, that modifies a specific DNA target sequence throughout the host genome and thereby defines self versus non-self (21,22); (iii) PglZ, which exhibits nuclease activity, (iv) PglZ’s tightly associated binding partner BrxB, which contains an inactivated AAA+ ATPase domain (23); (v) BrxA, a small DNA binding protein of unknown recognition specificity (24); and (vi) the DNA-binding AAA+ ATPase BrxL (25), which is replaced by various helicase enzymes in some non-Type I BREX systems (15).

**Figure 1.**
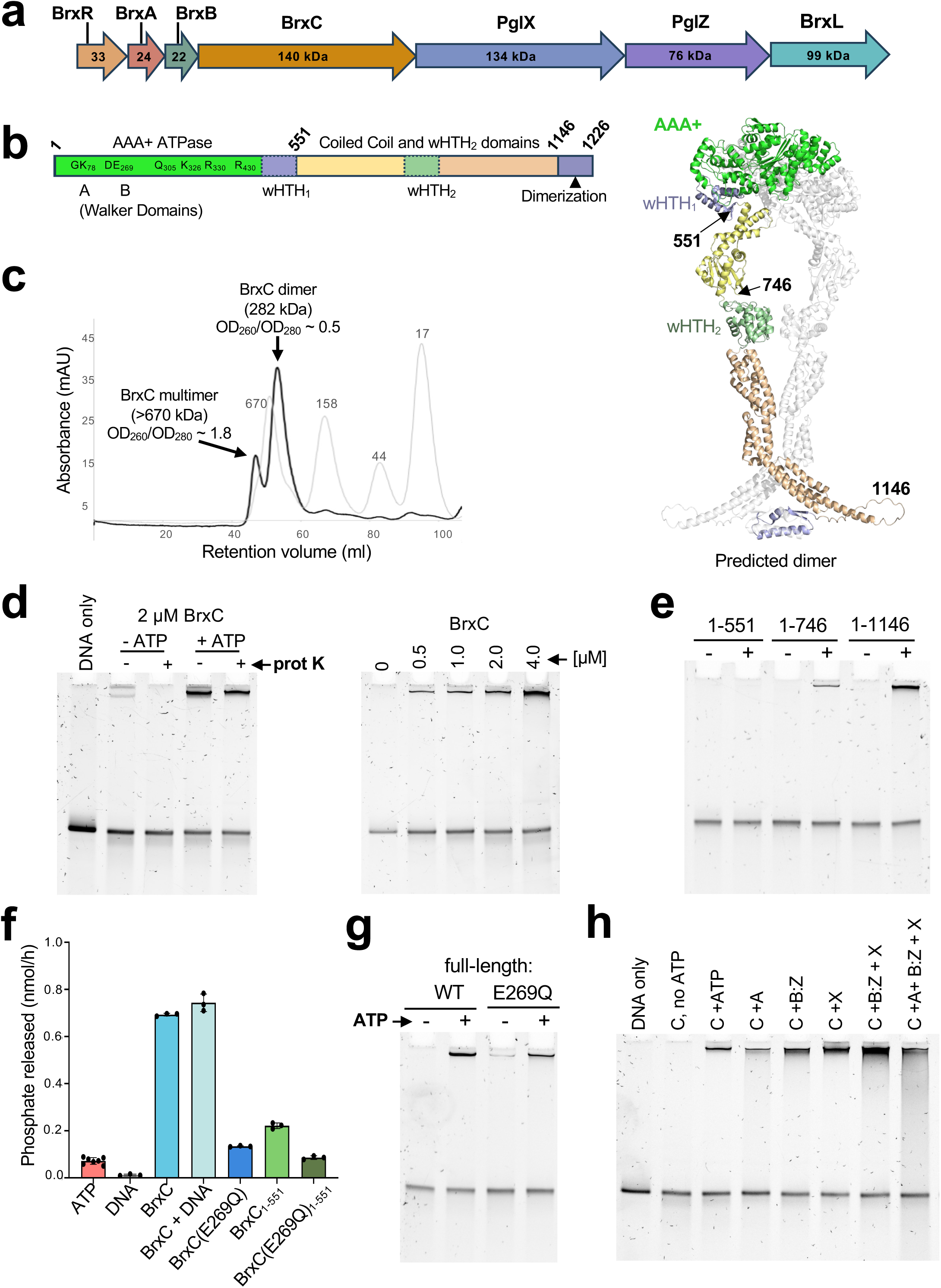
BrxC from *Acinetobacter* is a predicted dimer that binds DNA in an ATP-dependent manner. ***Panel a:*** The BREX system from *Acinetobacter* (BREX^Aci^) comprises a single operon encoding seven proteins. ***Panel b:*** BrxC is a multidomain protein of approximately 140 kDa. The N-terminal region (residues 1–551) comprises an AAA+ ATPase domain with conserved Walker A and B motifs and basic residues that promote ATP-dependent multimerization, together with a wHTH domain. The remainder of the protein comprises a central region (residues 551–1146) predicted to adopt an extended architecture that includes a second wHTH domain, followed by a C-terminal dimerization domain (residues 1146–1226). AF3 (32) predicts that BrxC^Aci^ forms a dimer (ipTM 0.37; pTM 0.40), with dimerization mediated by contacts between the AAA+ domains and C-terminal region of each monomer. The amino acid positions of truncations used in this study are indicated. ***Panel c:*** Size exclusion chromatographic analysis of full-length BrxC^Aci^, after an initial affinity tag purification step. BrxC^Aci^ elutes in two distinct peaks: an early-eluting peak associated with significant bound nucleic acid (OD260/280 ∼1.8),and a late-eluting peak corresponding to a BrxC^Aci^ dimer (∼280 kDa) with a much lower OD260/280 (∼0.5). The elution volumes of protein standard are indicated. ***Panel d:*** *Left gel:* Electrophoretic mobility shift analyses (EMSA; gel shifts) of purified, DNA-free BrxC^Aci^ dimer indicates binding to a 100 bp dsDNA duplex in an ATP-dependent manner. The resulting DNA complex is resistant to proteinase K treatment. *Right gel:* Titration of BrxC^Aci^ at the indicated concentrations in the presence of 1 mM ATP shows concentration-dependent formation of the shifted DNA band. ***Panel e:*** EMSA assays with 2 µM final concentrations of the indicated BrxC^Aci^ truncations were incubated with and without ATP. ***Panel f:*** ATPase assays of the indicated BrxC^Aci^ constructs, each incubated at 1 µM concentration. Where indicated, 100 bp dsDNA was added at 50 nM final concentration. ***Panel g:*** EMSA assays of full-length BrxC^Aci^ and BrxC^Aci^(E269Q), each at a final concentration of 2 µM, in the presence and absence of ATP. ***Panel h:*** EMSA assays of full-length BrxC^Aci^ in the presence of the indicated BREX^Aci^ proteins, all at 2 µM final concentration. All reactions included ATP except lane 2.

The final BREX factor, BrxC, is a large protein (>130 kDa) with an N-terminal AAA+ ATPase domain, a central region spanning two predicted winged helix-turn-helix (wHTH) domains as well as a coiled-coil domain, and a predicted C-terminal dimerization motif comprising the final 80 residues. (**Figure 1b**). Deletion of BrxC from an otherwise intact BREX system greatly reduces restriction activity; conversely BrxC expression is strongly upregulated when the *Acinetobacter* BREX system is derepressed via deletion of BrxR – a response that may mimic the induction of BREX activity that occurs during phage infection (18). Despite evidence suggesting a central role for BrxC in both the protective and restrictive activities of the BREX system, its behavior and role in either aspect of BREX function, such as perhaps modulating the switch from an inactive to active restriction system, has not yet been systematically examined.

The respective gene requirements for methylation and restriction have been characterized for several type I BREX systems. Across all systems examined, BrxB, BrxC, PglX, and PglZ are required for both methylation and restriction, whereas BrxA is required in some systems but not others (18,21,26–28). BrxL is dispensable for methylation in all systems analyzed and displays variable importance in phage restriction (18,21,26–28). Consistent with these gene requirements, pull-down analyses have detected complexes containing BrxB, BrxC, PglX, and PglZ proteins (21,22), although stable assemblies of these factors have not been observed and remain uncharacterized. BrxB and PglZ form a particularly tight complex (‘B:Z’) in widely diverged BREX systems (23); given their conservation across BREX systems (17), a B:Z complex may serve as an “anchor” within larger BREX subcomplexes. The PglX methyltransferase recognizes and acts at 6-basepair non-palindromic target sequences, but its activity requires BrxB, BrxC and PglZ (and in some systems, BrxA) (18,21,26–28).

The mechanism of BREX phage restriction remains poorly understood and has been reported to not occur via wholesale degradation of the phage genome (26), as is observed in the presence of active restriction endonuclease or CRISPR nuclease systems. However, we recently showed that PglZ exhibits nuclease activity—detectable as both nicking and double-strand DNA cleavage (23). Together, these data support a model in which overlapping sets of BREX proteins assemble into a methylation-directed complex under normal phage-free growth conditions and then reconfigure into a restriction-directed complex upon phage infection (21,22,26).

In this study, we characterize the solution behavior and multimerization properties of BrxC from a type I BREX system encoded in *Acinetobacter* and further demonstrate the generalizability and reproducibility of those results using a separate BREX system encoded in *Escherichia fergusonii*. We assess how specific residues and structural regions within BrxC (as well as ATP), drive those behaviors and describe a high-resolution cryo-EM structure that defines the basis for its ATP-dependent self-association. We further demonstrate that BrxC can associate with double stranded DNA (and in doing so, forms higher-order multimers) and/or with the BrxB-PglZ complex. We then visualized the latter complex via additional X-ray crystallography and EM analyses, and demonstrate that single point mutations in either BrxC or in BrxB (which respectively abrogate BrxC self-association or BrxC association with BrxB), cause different effects on the ability of the *Acinetobacter* BREX system to carry out host methylation and/or phage restriction.

Based on our results, we propose that BrxC may play a central role in regulating BREX function, perhaps influencing the system’s switch from an idling restriction negative (“R-”) state in the absence of a phage challenge to a restriction active (“R+”) state when a phage challenge is sensed — for example, through a change of its expression level, self-association, and/or binding affinities to other BREX factors and DNA.

## METHODS

### Expression and purification of *Acinetobacter* BREX proteins

The constructs generated and used throughout this study are listed in **Supplementary Table S1**. Genes encoding the *Acinetobacter* BREX proteins were subcloned from the native operon into pET15b. BrxC and PglX were expressed with an N-terminal Twin-Strep tag containing a TEV protease cleavage site, while PglZ and BrxA were expressed with a C-terminal Twin-Strep tag containing a thrombin cleavage site. The gene encoding *Acinetobacter* BrxB was subcloned into pET24d+ with no affinity tag; BrxB was insoluble unless co-expressed alongside PglZ. *E. coli* BL21-CodonPlus (DE3)-RIL cells were transformed or co-transformed with the expression vectors and used for protein expression. For co-expression studies, pET15b and pET24d+ vectors containing various combinations of *Acinetobacter* BREX genes were co-transformed into *E. coli* and expressed as described below. In some cases, pairs of genes from the native *Acinetobacter* operon were subcloned into the same expression vector; examples include PglX and PglZ, and BrxB and BrxC. In co-expression experiments, only one gene contained a Twin-Strep tag for affinity purification.

For autoinductions, 1 L ZY media containing relevant antibiotics (0.1 mg/ml ampicillin for pET15b, 0.05 mg/ml kanamycin for pET24d) in 2.8 L flasks were inoculated with ten single-colonies and incubated for 9 hours at 37 °C followed by 24 hours at 18 °C shaking at 175 rpm. Cells were harvested by centrifugation at 7,800 x *g* for 10 min at 4 °C. Cell pellets were resuspended and washed in ice-cold Buffer W (200 mM NaCl, 100 mM Tris pH 8.0, 1 mM EDTA pH 8.0) and re-pelleted at 3,200 x *g* for 12 min at 4 °C and stored at −20 °C.

For IPTG inductions, one colony per 10 ml of media was used to inoculate LB with antibiotics for overnight growth at 37 °C with shaking at 175 rpm. The overnight cultures were diluted 1:100 to inoculate 1 L LB media containing relevant antibiotics in 2.8 L flasks. The outgrowths were incubated at 37 °C, shaking at 175 rpm until an OD_600_ between 0.6 and 0.8 was reached, then IPTG was added to a concentration of 0.5 mM, followed by 24 hr at 18 °C shaking at 175 rpm. Cells were harvested by centrifugation at 7,800 x *g* for 10 min at 4 °C. Cell pellets were harvested and stored as described above.

For purification, each cell pellet was resuspended in 25 ml Buffer W followed by the addition Triton X-100 to a final concentration of 0.1%. Resuspended cells were lysed by sonication on ice and centrifuged in an SS34 rotor for 25 min at 18,000 x *g* and the supernatant (soluble fraction) was filtered through 5 µm syringe-filter. The filtered soluble fraction was then passed directly over 1 ml column volume (CV) Streptactin-4Flow resin (IBA Life Sciences) via gravity filtration column (Bio-Rad), collected, then passed over the column an additional time. The column and resin were then washed with 50 column volumes (CV) of ice-cold Buffer W. Elutions were performed with six sequential additions of 0.75 CV Buffer E (IBA Life Sciences), each incubated for several min before collection. Biotinylated thrombin (EMD Millipore) was added to eluted protein (∼1 unit thrombin per mg of protein). For purifications containing BrxC or PglX, His6 tagged-TEV protease was used, as thrombin had off-target activity with both proteins. The protease tag removal step was omitted from TST-tagged BrxC constructs lacking the TEV-protease recognition motif. Removal of the TST tags was assessed via ‘pre-’ and ‘post-protease’ gels on 4-12% BOLT SDS PAGE gel in MES buffer (Invitrogen), while MOPS buffer (Invitrogen) was used with PglX and BrxC to achieve adequate separation for analysis. Samples were then concentrated to ∼2 ml in an Amicon Ultra centrifugal filter (10,000 and 30,000 MWCO; Millipore), filtered over a 0.22 µm spin filter and loaded onto a HiLoad 16/600 Superdex 200 prep grade size exclusion column (Millipore Sigma) equilibrated in 25 mM Tris pH 7.5, 200 mM NaCl. Peak fractions were pooled and concentrated to a final concentration (following addition of 10% glycerol) of 44 µM for all BREX proteins or constructs. Single use aliquots were flash frozen in liquid nitrogen and stored at −80 °C.

Purification of co-expressed proteins was performed using the same procedure described above. In these experiments, one protein carried a twin-Strep tag, while the co-expressed partner proteins were untagged.

### Expression and purification of *Escherichia fergusonii* BREX proteins

For large scale expression of *E. fergusonii* BREX proteins, *E. coli* Rosetta (DE3) pLysS were transformed with pTRB791 (BrxC^Eferg^) and pTRB792 (BrxC^Eferg^(E268Q)). *E. coli* ArcticExpress (DE3) (Agilent) were transformed with pTRB789 (BrxC^Eferg^_1-551_) and pTRB790 (BrxC^Eferg^(E268Q)_1-551_). All proteins were expressed with an N-terminal His_6_-SUMO tag containing a human sentrin/SUMO-specific protease 2 (hSENP2) cleavage site. BrxB and PglZ were expressed and purified as described previously (23).

For autoinductions, single colonies were used to inoculate 150 ml of LB with 0.1 mg/ml ampicillin and 0.025 mg/ml chloramphenicol for overnight growth at 37 °C shaking at 180 rpm. Starter cultures were re-seeded 1:100 (v/v) into 12x 2 L baffled flasks containing 1 L ZY media and the relevant antibiotics and incubated at 37 °C for 6 hr then at 18 °C for 24 hr shaking at 150 rpm.

For IPTG inductions, single colonies were used to inoculate 70 ml of 2x YT with 0.1 mg/ml ampicillin and 0.01 mg/ml tetracycline for overnight growth at 37 °C shaking at 180 rpm. Starter cultures were re-seeded 1:100 (v/v) into 6x 2 L baffled flasks containing 1 L 2x YT media and incubated at 37 °C shaking at 150 rpm until the OD_600_ reached approximately 0.6. The temperature was reduced to 12 °C, IPTG was added to a final concentration of 1 mM, and expression was carried out for 48 hr.

Cell pellets were harvested by centrifugation at 4000 x *g* for 30 min at 4 °C. Cell pellets were resuspended in ice-cold A500 (500 mM NaCl, 20 mM Tris HCl pH 7.9, 10 mM imidazole, 10% v/v glycerol). Resuspended pellets were disrupted by sonication on ice (45% amplitude, 10 sec on, 20 sec off pulse intervals, 3 min) and clarified by centrifugation at 48,000 x *g* for 1 hr at 4 °C. Clarified cell lysates were loaded onto a 5 ml HisTrap HP column (Cytiva) pre-equilibrated with A100 (100 mM NaCl, 20 mM Tris HCl pH 7.9, 10 mM imidazole, 10% v/v glycerol). The HisTrap column was then washed with 50 ml A100.

For truncated BrxC proteins (BrxC^Eferg^_1-551_ and BrxC^Eferg^(E268Q)_1-551_), proteins were eluted directly onto a pre-equilibrated 5 ml HiTrap Q HP column using B100 (100 mM NaCl, 20 mM Tris HCl pH 7.9, 250 mM imidazole, 10% v/v glycerol). The Q HP column was washed with 50 ml A100 and transferred to an Åkta^™^ Pure (Cytiva), and the target protein was eluted by anion exchange chromatography using a salt gradient from 100% A100 to 60% C1000 (1 M NaCl, 20 mM Tris HCl pH 7.9, 10% v/v glycerol). Chromatographic peak fractions were collected, pooled, and incubated overnight in the presence of hSENP2 to facilitate the cleavage of the His-SUMO tag at 4 °C. For full-length BrxC proteins (BrxC^Eferg^ and BrxC^Eferg^(E268Q)), proteins were eluted in B100 and dialysed overnight into 5 L A100 using Pur-A-Lyzer™ dialysis tubing with a MW cutoff of 6 kDa (Sigma Aldrich) with gentle stirring at 4 °C. The following day, the hSENP2-treated samples were applied to a second HisTrap HP column pre-equilibrated in A100. The flow-through containing untagged target protein was collected and concentrated by centrifugation using the appropriate MWCO Vivaspin concentrator (Sartorius). Concentrated truncated BrxC^Eferg^ protein samples were applied to a HiPrep^™^ 16/60 Sephacryl^®^ S-200 HR column (Cytiva) and full length BrxC^Eferg^ protein samples were applied to a HiPrep^™^ 16/60 Sephacryl^®^ S-300 HR column (Cytiva), both pre-equilibrated with 1.2 column volumes (CV) of sizing buffer (500 mM NaCl, 50 mM Tris HCl pH 7.9, 10% v/v glycerol), for further purification by size exclusion chromatography (SEC). SEC peak fractions were pooled and analyzed by SDS-PAGE, then concentrated as described previously and quantified using a NanoDrop 2000 Spectrophotometer (Thermo Fisher). Final purified samples for biochemical analysis were resuspended in a 1:2 mixture of protein sample:storage buffer (500 mM NaCl, 50 mM Tris HCl pH 7.9, 70% v/v glycerol) and flash frozen in liquid nitrogen for storage at −80 °C.

### Mass photometry

Mass photometry experiments were performed for both BREX systems on a Refeyn TwoMP instrument, using the AcquireMP 2024 R2 and DiscoverMP v2024 R2 for data acquisition and analysis, respectively. DiscoverMP v2024 R2 was used to create figures. For each sample to be tested, the droplet dilution autofocus function was used to find the focus plane using 18-19 µl of buffer (200 mM NaCl, 25 mM Tris) for *Acinetobacter* proteins, or 9 µL PBS for *E. fergusonii* proteins, on uncoated glass slides (Refeyn). 1 or 2 µl of freshly prepared 100 nM protein sample was diluted directly into the drops (to reach final concentrations ranging from 5-10 nM) and 1 min video measurements were collected. *E. fergusonii* proteins were pre-incubated in 1x ZB (150 mM NaCl, 50 mM Tris HCl pH 8) with 10 µM MgCl_2_ in the presence and absence of 1 µM ATP for 1 hr at 37 °C. Experiments containing nucleic acids or protein:nucleic acid complexes were performed on glass slides (Refeyn) coated with poly-L-lysine (PLL). To prepare PLL-coated slides, 7 ul of 0.01% PLL (Sigma-Aldrich) was incubated between two coverslips for 30 sec, then separated and immersed in MilliQ water to remove excess PLL. The treated surface was then rinsed directly with MiliQ water from a squirt bottle and dried with clean, pressurized air. MassFerence P1 Calibrant (Refeyn) standards were used according to manufacturer specifications. Gaussian fits were used for measurements unless otherwise specified.

### DNA binding

A 100 bp dsDNA containing a PglX^Aci^ recognition target sequence (5ʹ-GTAGAT-3ʹ; **Supplementary Table S1**) was used as the substrate in EMSA assays with BrxC^Aci^. The dsDNA template was ordered from Integrated DNA Technologies (IDT) as a pre-annealed, HPLC-purified product. 20 ng of DNA was used in binding reactions. Untagged BrxC^Aci^ protein was diluted in 150 mM NaCl, 25 mM Tris (pH 7.5), and used at final concentrations of 0.5 - 4 µM. Binding reactions (20 µl) were assembled in binding buffer (50 mM Tris, pH 7.5; 50 mM NaCl, 1 mM DTT, 5 mM MgCl2). When included, ATP was added at a final concentration of 1 mM. Reactions were incubated at 30 °C for 2 hr. Where indicated, 1 unit of proteinase K was added per 20 µl reaction and incubated for 30 min at 42 °C. Samples were resolved on native 8% acrylamide gels (prepared using 29:1 acrylamide:bisacrylamide; Bio-Rad) in 0.5x TBE buffer. 5 µl of each binding reaction were loaded on the gel and run with 1x TBE running buffer at room temperature for 30 min at 150 V. Gels were stained for 30 min in SYBR Gold (Invitrogen) diluted in 1x TBE, rinsed twice with water, and imaged on a BioRad Gel Doc XR+ scanner.

### Protein-protein association pulldown assays

For pulldown assays using purified proteins from *Acinetobacter*, the protein isolated by affinity purification carried a twin-Strep tag, whereas all partner proteins were either untagged or contained a His₆-tag. The Twin-Strep-tagged protein was incubated at a 2-fold molar excess (final concentration 2 µM) relative to the partner proteins (final concentration 1 µM) in binding buffer (150 mM NaCl, 25 mM Tris, pH 7.5) in 50 µl reaction volumes for 25 min at 22 °C. Streptactin agarose beads (IBA Lifesciences Inc., cat # 2-1250-002) were prepared by pipetting 12 µl of a 50% slurry into a 0.6 ml Eppendorf tube and centrifuged at 2000 x g for 1 min. The supernatant was removed by aspiration using a 25-gauge needle attached to a vacuum apparatus and 100 µl of water was added. This was repeated more with binding buffer. Final supernatant was removed by aspiration and 50 µl of binding buffer was added to resuspend the beads.

After the protein incubation reactions were complete, samples were added to the Streptactin beads, gently tapped to resuspend, and incubated on a LabQuake rotisserie at 4 °C for 45 min, with beads resuspended by tapping every 10 min. The beads were centrifuged as above, a sample of the supernatant was collected (“flow-through”), and the remaining supernatant was removed by aspiration. The beads were then washed twice with binding buffer using the same procedure described above to prepare the beads, transferred to a fresh tube, and washed two additional times. After the final wash and aspiration, 40 µl of Buffer E elution buffer (X) was added and incubated at 22 °C for 15 min. Samples were centrifuged for 1 min at 5000 x g, and 10 µl of the supernatant (“elution”) was removed without disturbing the beads. An equal volume of 2x BOLT SDS sample buffer (containing 100 mM DTT) was added. Samples were boiled and resolved on 4–12% BOLT gels (Invitrogen) using MES running buffer for 25 min at 200 V. Gels were visualized by Coomassie blue staining or silver staining.

Purification of co-expressed proteins was carried out using the same procedure described above for individually expressed proteins.

### Analytical size exclusion chromatography

Analytical SEC was performed on an Åkta™ Pure FPLC system (Cytiva). Protein samples were made up to 10 µM in A-SEC buffer (150 mM NaCl, 20 mM Tris HCl pH 7.9) to a final volume of 100 µl and incubated either at room temperature (BrxC^Eferg^ and BrxC^Eferg^(E268Q) or at 37 °C (BrxC^Eferg^_1-551_ and BrxC^Eferg^(E268Q)_1-551_) for 1 hr with and without 1 mM ATP and 10 mM MgCl_2_. Samples were loaded onto a 100 µl capillary loop using a 100 µl Hamilton syringe. The loop was washed with 500 µl nuclease-free water followed by 500 µl A-SEC buffer before and between each run. Samples were loaded onto a Superdex™ 200 increase 10/300 GL SEC column (Cytiva), pre-equilibrated with 1.2 CV A-SEC buffer, by running A-SEC buffer through the loop at 0.375 ml/min. Samples were resolved across 1.2 CV. In cases where the content of chromatogram peaks required verification by SDS-PAGE, 0.5 ml fractionation was performed in 96-well deep-plate blocks.

### ATPase activity assays

ATPase activity assays with *Acinetobacter* and *E. fergusonii* BREX proteins were performed with BIOMOL® Green (Enzo Life Sciences) in a 96-well plate format. A standard curve was generated from serial dilutions of free phosphate and used to calculate phosphate release from experimental samples. Experimental wells containing the protein of interest were set up in 50 μl total volumes; A-SEC buffer was used for BrxC^Eferg^ assays, while a buffer containing 150 mM NaCl and 20 mM Tris pH 7.5 was used for BrxC^Aci^ assays. For reactions with BrxC^Eferg^ and BrxC^Aci^ constructs alone, proteins were incubated at 500 nM, 1 μM, and 2 μM. For complexing assays, BrxC^Eferg^_1-551_ and BrxC^Eferg^(E268Q)_1-551_ were incubated at 1 μM with PglZ^Eferg^ and/or BrxB^Eferg^ at 1 μM and 0.5 μM. BrxC^Aci^ constructs (including truncations and point mutants) were incubated at 1 μM with PglZ^Aci^/BrxB^Aci^ at 1 μM. For assays containing DNA, a 100 bp dsDNA containing a PglX^Aci^ recognition sequence (**Supplementary Table 1**) was added to a concentration of 50 nM and incubated with proteins at 1 μM each. Reactions were carried out with and without 100 μM ATP and 10 mM MgCl_2_ for 1 hr at 37 °C. Negative control wells contained ATP or protein only. Positive control wells contained 1 U FastAP thermosensitive alkaline phosphatase (Thermo Fisher Scientific). Reactions were terminated by the addition of 100 μl BIOMOL green reagent and the color was allowed to develop for 15 min. Absorbance was measured in 96-well format at 620 nm. Readings were blanked using BIOMOL green reagent and interpolated on a standard curve to give the amount of free phosphate produced. Values were averaged from three independent wells.

### Cryogenic-Election Microscopy (cryo-EM) analyses

Flow charts and summaries of data collection and processing methods and corresponding structural modeling, as described below, are shown in **Supplementary Figure S1 and Supplementary Table S2**.

#### BrxC^Aci^_E269Q_ negative stain screening

Negative stain grids were prepared by applying 3 μl of SEC purified sample at a concentration of approximately 0.02 mg/ml to a plasma-cleaned Formvar/Carbon 400 mesh Copper grid (Ted Pella). The sample was allowed to absorb for 30 sec before wicking excess solution with filter paper. The grid was quickly washed two times in 30 μl drops of water and once in a 30 μl drop of 2% uranyl formate (UF) followed by a final staining for 60 sec with another 30 μl drop of 2% UF. The grids were air dried for at least 1 hr. Grids were screened on a Talos L120C transmission electron microscope (Thermo Fisher), operating at 120 kV and equipped with a 4k x 4k Ceta CMOS high-resolution 16M camera (Thermo Fisher). The sample distributed homogeneously and at random orientations over the surface of the prepared negative stained grids.

#### BrxC^Aci^_E269Q_ data collection and processing (Supplementary Figure S1a)

C-Flat 1.2/1.3 holey carbon film coated 300 mesh copper grids (Electron Microscopy Sciences) were prepared for cryo-EM by plasma cleaning with water vapor using a Tergo EM Plasma Cleaner (Pie Scientific). 3 μl of SEC purified sample at a concentration of 0.4 mg/ml was applied to the prepared grids, which were then blotted for 5.0 sec at a tension of 0 and plunge-frozen into liquid ethane using a Mark IV Vitrobot (Thermo Fisher). A dataset of 4131 movies was collected at a super resolution pixel size of 0.56 Å using a Glacios 200 kV electron microscope (Thermo Fisher) equipped with a Gatan K3 direct electron detector. Movies were imported into CryoSPARC (29), motion and CTF corrected, denoised using 100 training micrographs, and particles were picked by automated searching for Gaussian signals on 200 micrographs. This resulted in approximately 100,000 particles which were extracted to a box size of 360 pixels (px) and Fourier cropped to 180 px and classified into 50 2D classes. Eleven classes containing approximately 40,000 particles with the best alignments were selected and passed into an Ab-initio 3D reconstruction job with 4 classes followed by further Heterogenous Refinement (Hetero refinement), resulting in three classes containing the majority of particles and a single junk class containing 3 particles. The particles from the three non-junk classes were again sorted into 50 2D classes and the best alignments were chosen as templates for picking on the entire dataset. Approximately 1.77 million particles were extracted into a box size of 360 px and fed into a Hetero refinement job using initial volumes (3 unique, 1 junk) from the particle subset Hetero refinement, resulting in 2 unique volumes and 2 junk volumes.

The 2 unique volumes (subsets A and B with 460k and 550K particles, respectively) were further individually refined using Non-Uniform Refinement (NU refinement) (30). The A and B NU refinement volumes were individually subjugated to Hetero refinement, with input volumes for volume A being two copies of the NU refinement volume and for volume B being the NU refinement volume plus a junk volume. This resulted in two unique volumes from subset A and a third unique volume from subset B. All three unique volumes with their associated particles plus a junk volume were combined into a final Hetero refinement, resulting in 3 unique volumes which were individually run through NU refinement.

One volume required the hand to be flipped, then the three volumes were aligned to allow for easier comparison. Two of the three volumes were run through Global and Local CTF refinement, the third volume did not benefit from these refinements. A final round of NU refinement was performed resulting in three volumes with 477k, 315k, and 191k particles, and GSFSC resolutions of 2.95Å, 3.02 Å, and 3.37Å, respectively.

#### Model fitting (Supplementary Figure S1b)

Model fitting was primarily accomplished with the 2.95 Å volume. The volume was loaded into ChimeraX (31) along with a dimeric AlphaFold3 (32) model of BrxC^Aci^ extending from residues 1 to 1225 containing ATP. The model was roughly placed in the volume with the Fit in Map function, then refined with minor rebuilding using ISOLDE (33) and Coot (34). Model fitting for the 3.02 Å and 3.37 Å volumes primarily used the model fit in the 2.95 Å volume and supplemented with the AlphaFold model for residues not present in the 2.95 Å resolution volume. Those models were also refined with minor rebuilding using ISOLDE and Coot.

#### Visualization of Acinetobacter BrxC-B:PglZ complexes (Supplementary Figures S1c, d, and e)

The same general methods as BrxC^Aci^(E269Q) were followed for samples of a complex of the BrxC^Aci^ N-terminal domain (residues 1 to 553) bound to BrxB^Aci^ and PglZ^Aci^ with minor differences noted here. Stable particles containing all three subunits were generated by fusing the C-terminal end of BrxC^Aci^ to the N-terminal end of BrxB^Aci^ (a construct that was enabled by computational modeling using AlphaFold3 (32) in conjunction with crystallographic analyses described below) and then adding purified PglZ to that construct in a 1:1 molar ratio. Grids and corresponding data that led to visualization of a monomeric *Acinetobacter* C-B:Z complex contained no ATP or MgCl_2_ in the sample buffer. Grids and corresponding data that led to visualization of a dimeric *Acinetobacter* (C-B:Z)_2_ complex corresponded to a higher protein concentration (3.5 mg/mL) in the presence of 1 mM AMP-PNP, 1 mM MgCl_2_, and 0.05% CHAPS. Initial full model fitting was performed with AlphaFold3 models in ChimeraX using the ‘Fit in Map’ tool. Individual domains were then further fit into volumes and minimally refined with ISOLDE to resolve distortions arising from manual fitting. Low resolution and lack of detail prevented further rebuilding and refinement.

### X-ray crystallography

Sequences encoding BrxC^Aci^ (E269Q)_1-551_ and PglZ^Aci^_1-98_ were subcloned into pET-based expression vectors containing protease-cleavable Twin-Strep affinity tags; wild-type BrxB^Aci^ was subcloned without affinity tags. Wild-type BrxB^Aci^ and strep-tagged PglZ^Aci^_1-98_ were co-expressed and co-purified, pulling down on PglZ^Aci^_1-98_, while BrxC^Aci^(E269Q)_1-551_ was expressed and purified separately. Affinity tags were removed by protease digest and further purified as described above prior to setting up crystallization trials. BrxC^Aci^(E269Q)_1-551_ and the PglZ^Aci^_1-98_:BrxB^Aci^ complex were incubated prior to crystallization for 30 min at room temperature in a 1:1 molar ratio in buffer with 1 mM ATP and 10 mM MgCl_2_ added. Crystals were grown by hanging drop vapor diffusion in drops set with 1 µl of protein (protein concentration 9 mg/ml, 94 µM for each individual component) plus 1 µl of well solution (250 mM NaCl, 100 mM HEPES pH 8.5, 21% PEG3350). For cryopreservation, cryoprotectant (300 mM NaCl, 100 mM HEPES pH 8.5, 25% PEG3350, 25% 2-Methyl-2,4-pentanediol) was added to drops containing crystals through sequential additions and equilibrated for 1 min before flash freezing with liquid nitrogen.

The crystals were found to display I2 crystallographic symmetry, with unit cell dimensions a = 149.2 Å, b = 115.0 Å, c = 194.1 Å, and β = 108.0°. Data collection was conducted at beamline 5.0.1 the Advanced Light Source (ALS) synchrotron beamline 5.0.1 at the Lawrence Berkeley National Laboratory (LBNL) in Berkeley California. Data was collected on a Pilatus area detector and processing using program XDS. The X-ray diffraction data (**Supplementary Table S3**) was integrated and scaled to 2.74 Å resolution, corresponding to an I/σ(I) cutoff of 2.0. The structure was solved via molecular replacement (Top LLG=2412; Top TFZ = 21), using the refined model of the BrxC^Aci^_1-551_ dimer solved by cryo-EM and an AlphaFold3 model of the BrxB^Aci^:PglZ^Aci^_1-98_ complex as sequential search models. MR produced a solution corresponding to an asymmetric unit containing two copies of a complex of *Acinetobacter* (BrxC_1-551_)_2_BrxB:PglZ_1-98_. The final values for R_work_/R_free_ were 0.195 / 0.250.

### Phage Restriction Assays

For assays testing the *Acinetobacter* BREX system, bacteriophage *λ*_JL801_ was obtained from the Gerald Smith Lab at Fred Hutchinson Cancer Center. Serial dilutions of λ_JL801_ were prepared in phage buffer (50 mM Tris pH 7.5, 25 mM NaCl, 4 mM MgSO_4_). *E. coli* (NEB ER2683) was transformed with pACYC184 (“Empty vector”), pACYC BREX WT, pACYC BREX #x0394;BrxC, pACYC BREX ΔBrxB, pACYC BREX BrxC(E269Q), pACYC BREX BrxC(R330E), pACYC BREX BrxC(1–1146), pACYC BREX BrxC(1–551), or pACYC BREX BrxB(R82E) and grown overnight. Overnight cultures were diluted 1:33 into 20 ml of fresh LB containing 1.25 mM MgCl_2_ and grown to OD_600_ between 0.35 and 0.45. 200 ul of the outgrowth cultures were pipetted into 4 ml of 0.5% molten LB agar containing 1.25 mM MgCl_2_ and poured onto 1.5% LB plates containing 1.25 mM MgCl_2_ and chloramphenicol (0.025 mg/ml). Once solidified, the plates were spotted with 5 µl 10-fold serial dilutions of λ_JL801_ and allowed to incubate overnight at 37 °C before plaque forming units (pfu/ml) were counted on each plate. Efficiency of Plating (EOP) values were calculated by determining the phage titer on a test strain divided by the titer on the control strain.

For assays testing the *E. fergusonii* BREX system, *E. coli* DH5α were transformed with pTRB563 (pBrxXL), pTRB564 (pBrxXL-Δ*pglX*), pTRB565 (pBrxXL-Δ*brxU*), pTRB566 (pBrxXL-Δ*brxU*Δ*pglX),* pTRB804 (pBrxXL-Δ*brxC*), pTRB805 (pBrxXL-*brxC_1-551_*), pTRB806 (pBrxXL-*brxC(E268Q)*), a vector-only control (pTRB507), and grown overnight. Serial dilutions of phage Pau (from the Durham Collection (27)) were produced in phage buffer (10 mM Tris HCl pH 7.4, 10 mM MgSO_4_, 0.01% (v/v) gelatin). 200 μl of overnight culture and 10 μl of phage dilution were added to a sterile 8 ml plastic bijoux with 3 ml of 0.35% (w/v) LB-agar and poured onto LB plates. Plates were incubated overnight at 37 °C before plaque forming units (pfu) were counted on each plate. EOP values were calculated by determining the phage titre on a test strain divided by the titre on a control strain.

For each construct tested, a minimum of three and up to five biological replicate experiments were performed, each of which corresponded to three technical replicates. Mean values and standard deviations were plotted in GraphPad Prism.

### Pacific biosciences sequencing

Libraries for methylation sequencing were prepared using the SMRTbell HiFi 96 Prep kit (Pacific Biosciences). *E. coli* strain ER2796 (35) was transformed with the respective constructs as for phage restriction assays above. Bacterial gDNA was extracted from 5 ml of overnight culture using a Monarch Genomic DNA Purification Kit (New England Biolabs #T3010). Bacterial gDNA samples were then sheared using a Qiagen Tissue Lyser II at 30 Hz for 240 sec to produce DNA fragments with a mean size of 8–10 kb. The DNA was damage- and end-repaired, and SMRT-bell adapters were then ligated. Exonuclease treatment removed non-incorporated SMRT-bell DNA. Sequencing was performed on a PacBio Revio (Pacific Biosciences). Data were analysed using PacBio SMRTAnalysis on SMRTLink_25.1 software Base Modification Analysis for Sequel data, to identify DNA modifications and their corresponding target motifs.

## RESULTS

### BrxC ATP-dependent assembly and DNA binding behavior

The results presented below primarily focus on BrxC and associated proteins from *Acinetobacter* (BrxC^Aci^), which are then supported by comparative experiments using BrxC from *E. fergusonii* (BrxC^Eferg^). The first step of this project was to purify the full-length BrxC^Aci^ and BrxC^Eferg^ homologs and assess their solution behaviors, self-assemblage states, and possible DNA binding activities.

#### BrxC purification and initial solution behavior

Full-length wild-type (WT) BrxC^Aci^ and BrxC^Eferg^ homologs were each expressed and purified from *E. coli* as described in **Methods**. Size exclusion chromatography (SEC) of both constructs revealed two sequentially eluting species (**Figure 1c and Supplementary Figure S2a**): an early arriving peak corresponding to >670 kDa total mass (assessed relative to the largest molecular weight marker) with a high OD_260_/OD_280_ ratio >1.5, consistent with a large multimeric protein complex co-eluting with bound nucleic acid; and a later arriving, higher amplitude, peak corresponding to a mass of slightly less than 300 kDa with a much lower OD_260_/OD_280_ ratio (∼0.5-0.6), consistent with a BrxC dimer containing little or no bound nucleic acid. Once purified and separated from the larger protein-DNA fractions, the smaller species maintained the same elution behavior and apparent mass in subsequent SEC runs.

#### BrxC DNA binding and ATPase activity

The SEC profile described above led us to hypothesize that full-length BrxC purifies from *E. coli* as both a nucleic-acid–free dimer and a larger, heterogeneous population of DNA-bound multimers. The presence of a pair of winged-helix-turn-helix (wHTH) domains in BrxC, which are commonly associated with DNA binding, further suggested that BrxC might interact with DNA. To test this possibility, we incubated the purified BrxC^Aci^ dimer with double-stranded DNA (**Supplementary Table S1**) in the presence or absence of ATP and assessed binding by electrophoretic mobility shift assays (EMSAs). This analysis demonstrated ATP-dependent formation of a highly shifted BrxC^Aci^-DNA complex at low micromolar protein concentrations (**Figure 1d; left)**. To examine the stability of the shifted BrxC^Aci^–DNA complexes, EMSA reactions were treated with proteinase K **(Figure 1d; left**). The highly shifted band persisted following protease treatment, consistent with formation of a protease-resistant complex. Titration of BrxC^Aci^ in EMSA reactions induced a concentration-dependent increase in the highly shifted DNA species; however, complete shifting of the DNA substrate was not observed, even at the highest protein concentrations tested (**Figure 1d; right**).

To further characterize BrxC’s DNA-binding properties, we generated and purified truncated BrxC^Aci^ constructs corresponding to predicted boundaries between domains (indicated in **Figure 1b**). These truncations included (1) residues 1-551 (BrxC^Aci^_1-551_), spanning the N-terminal AAA+ ATPase and the first wHTH domain; (2) residues 1-746 (BrxC^Aci^_1-746_), which further incorporates the subsequent predicted central α + β domain; and (3) residues 1-1146 (BrxC^Aci^_1-1146_), which includes all but the final 80 residues that are predicted to form a C-terminal protein dimerization interface. BrxC^Aci^_1-1146_ demonstrated similar DNA binding as compared to the full-length constructs, whereas BrxC^Aci^_1-746_ showed significantly reduced but observable binding. No binding was observed for the BrxC^Aci^_1-551_ construct (**Figure 1e**).

Given BrxC’s predicted N-terminal AAA+ ATPase domain, and wondering if such activity might influence DNA binding, we next measured the ATPase activity of both BrxC homologs. Full-length BrxC^Aci^ (WT) and BrxC^Eferg^ (WT) both exhibited low but comparable levels of ATPase activity (**Figure 1f and Supplementary Figure S3**). Incubation with DNA induced only a marginal increase in BrxC^Aci^ (WT) activity. The activity of truncated BrxC^Aci^_1-551_ and BrxC^Eferg^_1-551_ constructs was substantially reduced compared to their full-length counterparts, despite the presence of an intact AAA+ ATPase domain (**Figure 1f and Supplementary Figure S3**). We also generated and tested mutants containing an E->Q point mutation in the conserved ATPase Walker B motif (BrxC^Aci^(E269Q) and BrxC^Eferg^(E268Q). (Similar mutations in other AAA+ ATPases have been shown to allow ATP binding while strongly inhibiting hydrolysis (36)). In both full-length constructs, this mutation caused significantly decreased ATPase activity (although some residual activity above background remained), and the same mutations in the 1-551 constructs further suppressed activity (**Figure 1f and Supplementary Figure S3**).

With these results in hand, we next asked whether the Walker B mutation affected DNA binding. The full-length BrxC^Aci^(E269Q) mutant retained ATP-dependent DNA-binding in EMSA assays (**Figure 1g)**. SEC analyses of BrxC^Eferg^(E268Q) also indicated dsDNA binding activity, based on a similar elution profile as BrxC^Eferg^ WT (**Supplementary Figure S2b**).

These experiments collectively indicate that (1) BrxC binds double-strand DNA in an ATP-dependent manner, with ATP hydrolysis not required for binding, and (2) that DNA binding requires the N-terminal ATP binding domain and the first coiled-coil region up to amino acid 746.

#### Effect of additional BREX factors on Acinetobacter BrxC DNA binding

We next assessed how other purified BREX proteins, individually and in combination, affected BrxC^Aci^’s interaction with DNA. These included (1) BrxA^Aci^, (2) a tightly associated complex of BrxB^Aci^ and PglZ^Aci^ (“B:Z”), and (3) PglX^Aci^. Incubation of BrxC^Aci^ with equimolar amounts of B:Z or PglX^Aci^ individually did not alter DNA binding, and both factors induced a modest increase in the shifted band when titrated at higher molar ratios (**Figure 1h, Supplementary Figure S4a**). In contrast, BrxA^Aci^ reduced BrxC^Aci^ DNA binding. When B:Z and PglX^Aci^ were incubated together with BrxC^Aci^, an increase in the shifted band was observed relative to either factor alone, whereas inclusion of BrxA^Aci^ with both factors reduced formation of the shifted band (**Figure 1h**). In comparison to BrxC^Aci^, the B:Z complex and PglX induced only modest shifts when tested on their own, demonstrating the observed DNA binding behavior is driven by BrxC^Aci^ (**Supplementary Figure S4a**).

Based on BrxA^Aci^’s ability to inhibit BrxC^Aci^ DNA binding in EMSAs, we tested whether the two proteins physically interact using a pull-down assay with purified proteins (**Supplementary Figure S4b**). This analysis indicated a direct interaction between the two proteins. However, they did not co-elute on SEC, suggesting that the interaction may be low-affinity or require additional factors for stabilization. Together, the EMSA and protein interaction analyses may support a model in which BrxA modulates BrxC activity.

#### BrxC self-association and multimerization

We next carried out a series of mass photometry experiments to further examine BrxC protein-protein association and dimerization behavior (**Figure 2**). Three protein constructs were examined: full length BrxC^Aci^ (BrxC^Aci^_1-1226_), BrxC^Aci^_1-1146_ and BrxC^Aci^_1-551_. Each construct was analyzed in either the WT ATPase background or with an inactivating E269Q mutation in the Walker B motif (six constructs total), all in the presence or absence of ATP.

**Figure 2.**
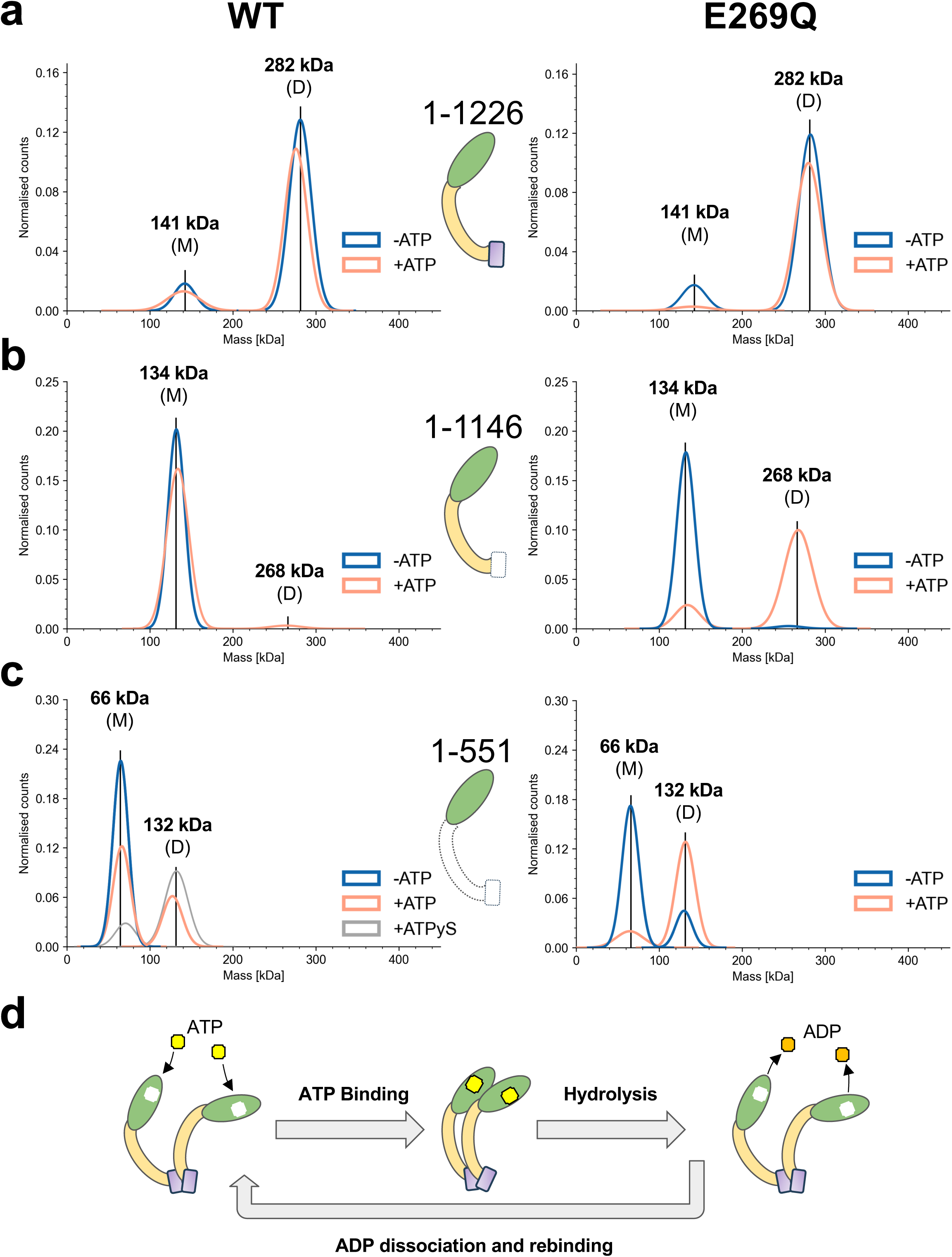
Solution behavior and ATP dependence of purified *Acinetobacter* BrxC constructs. Mass photometry was used to analyze the mass and multimerization behavior of full-length BrxC^Aci^ (BrxC^Aci^_1-1226_), BrxC^Aci^_1-1146_ and BrxC^Aci^_1-551_. Each construct was examined in the absence and presence of the inactivating E269Q point mutation in the ATPase Walker B motif (left plots and right plots, respectively) and in the absence and presence of ATP (purple and orange traces in each plot, respectively). ***Panel a:*** Full-length BrxC^Aci^ behaves largely as a dimer regardless of the presence or absence of ATP, both in its wild-type (WT) form (left) and with the E269Q mutation (right). ***Panel b:*** Deletion of the C-terminal region (residues 1147 to 1226) largely eliminates dimerization of the WT protein (left). Incorporation of the Walker B E269Q mutation rescues dimerization when ATP is present (right). ***Panel c:*** The N-terminal AAA+ domain (residues 1-551) is largely monomeric, with a measurable increase of the dimer in the presence of ATP (left). The fraction of ATP-dependent dimer is significantly increased by the Walker B E269Q mutation (right). ***Panel d:*** These analyses indicate that dimerization of BrxC is promoted both by its C-terminal region (residues 1147 to 1226) and by ATP-driven association of its N-terminal AAA+ ATPase domain. The latter association is further stabilized by the inactivating Walker B mutation (E269Q).

Full-length BrxC^Aci^ (WT) displayed a predominant mass of 282 kDa, corresponding to a protein dimer, and a minor monomeric species (141 kDa) regardless of the presence or absence of ATP (**Figure 2a, left**). In contrast, the same full-length construct harboring the E269Q mutation displayed a nearly identical distribution in the absence of ATP, but in the presence of ATP was predominantly dimeric, with negligible monomeric BrxC^Aci^ observed (**Figure 2a, right**).

Removal of the protein’s final 80 residues, generating BrxC^Aci^_1-1146_, resulted in a primarily monomeric species (134 kDa) in the presence or absence of ATP (**Figure 2b, left**), indicating that the final 80 residues of the protein mediate dimerization. When the catalytic E→Q mutation in the Walker B motif was incorporated into this truncated construct (BrxC^Aci^(E269Q)_1-1146_), the protein remained monomeric in the absence of ATP; however, in the presence of ATP, a significant fraction shifted to 268 kDa, consistent with a dimeric species (**Figure 2b, right**).

Finally, BrxC^Aci^_1-551_ was examined. In the absence of ATP, BrxC^Aci^_1-551_ **(Figure 2c, left)** was entirely monomeric (66 kDa), whereas a small but measurable fraction dimerized in the presence of ATP. In contrast, for BrxC^Aci^(E269Q)_1-551_, a minor dimeric species was observed without ATP but shifted mostly to the dimeric state in the presence of ATP (**Figure 2c**, **right**). We also tested the effect of the non-hydrolyzable analog ATPγS on BrxC^Aci^_1-551_ dimerization. In the presence of ATPγS, the majority of WT BrxC^Aci^_1-551_ was dimeric, phenocopying the E269Q mutant incubated with ATP (**Figure 2c, left**).

Similar analyses with *E. fergusonii* BrxC constructs, utilizing analytical-SEC and mass photometry, largely reproduced the results described above (**Supplementary Figure S5)**. Notably, as with BrxC^Aci^, full-length BrxC^Eferg^ exists primarily as a dimer; and the N-terminal ATPase domain in isolation (BrxC^Eferg^_1-551_) exhibits ATP-dependent dimerization.

These results collectively suggest that BrxC contains two regions that independently promote dimerization: the protein’s N-terminal AAA+ ATPase domain, which undergoes reversible homodimerization that is regulated by ATP binding and hydrolysis, and its final ∼80 residues (residues 1147–1226), which forms a stable, ATP-independent dimerization interface. These observations suggest a model (**Figure 2d**) in which BrxC dimerization is driven by two different regions of the protein and corresponding interactions: reversible ATP-binding and hydrolysis via the N-terminal ATPase domain, operating in concert with a constitutive dimerization motif located at the protein’s C-terminal end.

### A cryo-EM structure of the *Acinetobacter* BrxC dimer illustrates the involvement of ATP in protein-protein association

To determine the atomic basis for the described behaviors (**Figures 1 and 2**) we screened full-length BrxC^Aci^(E269Q) in the presence of ATP to produce uniform single-molecule particles suitable for visualization by cryogenic electron microscopy (cryo-EM). Our analysis (**Supplementary Figure S1a** and **S1b**) revealed a dominant species corresponding to a dimer of BrxC^Aci^’s N-terminal domains. Electron density for this region was uniformly well resolved, with an average resolution of ∼2.9 Å. The structure showed the N-terminal domains arranged in an asymmetric ‘back-to-face’ configuration, with ATP and coordinated magnesium ion positioned at the dimer interface and in the ATP binding pocket of the open-faced protomer (**Figure 3a**). BrxC^Aci^ coordinates ATP using conserved AAA+ ATPase residues, including K78 (Walker A), E269 (Walker B), and R430 (Sensor 2) from one BrxC^Aci^ subunit; and K326 and R330 (arginine fingers) from the partner subunit (**Figure 3b, left**). The residues involved in ATP binding listed above are well conserved across multiple BrxC species, including those examined as part of this study (**Supplementary Figure S6**) (36). Analysis of this structure reveals a potential mode in which additional BrxC subunits could stack onto the BrxC dimer, mediated by interactions between the Arg finger residues of the incoming subunit and the ATP pocket of its binding partner (**Figure 3b, right**).

**Figure 3.**
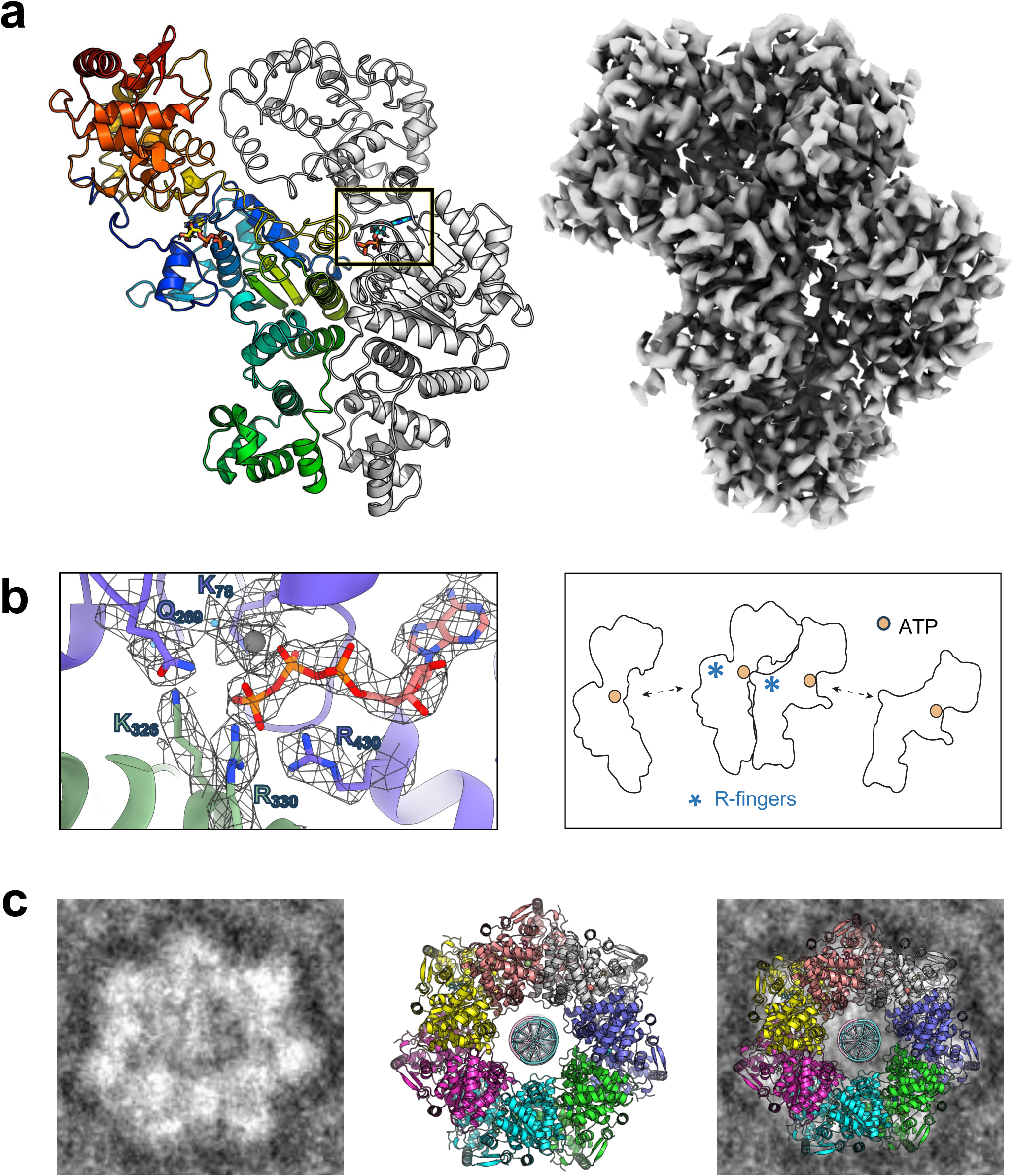
Cryo-EM analyses of the purified *Acinetobacter* BrxC dimer. ***Panel a:*** Full-length, purified BrxC^Aci^(E269Q) was examined via negative stain and subsequent cryo-EM. The resulting map (right) provided clear detail, to an average of approximately ∼2.9 Å resolution, for the N-terminal, ATP-bound domains (residues 1 to 551). At lower contour levels (**Supplementary Figure 1**), additional density corresponding to BrxC^Aci^’s central region extending beyond the AAA+ domain was observable. In the cartoon image (left) one subunit is colored as a spectrum from its blue N-terminus to its red C-terminus; the second subunit is shown as a light grey ribbon outline. ***Panel b, left:*** the indicated residues from each BrxC^Aci^ monomer interact with the bound ATP molecule and contribute to dimer formation. ***Panel b, right:*** BrxC^Aci^ monomers bind in a ‘back to face’ packing arrangement to form a dimer, with Arg finger residues K326 and R330 from one subunit making contacts to the ATP bound in its partner subunit. Additional BrxC monomers could bind to BrxC dimers as depicted in the cartoon. ***Panel c:*** Small numbers of particles observed throughout individual negative stain EM micrographs indicated the occasional presence of larger, well-ordered circular species displaying heptameric geometry. Predictive structural modeling of BrxC^Aci^ multimers using AlphaFold 3 produced circular, closed heptameric rings with dimensions that closely matched the observed particles, with an interior pore of sufficient diameter to accommodate a DNA duplex.

Whereas the electron density surrounding BrxC^Aci^’s N-terminal domains was well defined, the remainder of the full-length protein was largely disordered, with the exception of much lower resolution density features extending part way into the protein’s central region (predicted to form mostly helical coiled-coil structural motifs and the associated wHTH domain) that were visible only at significantly lower contour levels (**Supplementary Figure S1a, bottom)**. It is likely that a combination of preferred orientation of the N-terminal domains on the grid and significant conformational heterogeneity extending beyond that region resulted in the lack of clear density for the remainder of the protein structure.

Close examination of negative stain grids also indicated the infrequent presence of higher order circular protein multimers with seven-fold (heptameric) symmetry (**Figure 3c**). The shape and dimensions of those particles correspond closely to an AlphaFold3 model with high relative confidence (ipTM: 0.55, pTM 0.60, compared to the BrxC^Aci^ ATP-bound dimer predicted model with ipTM and pTM of 0.37 and 0.4 respectively) predicting a circular ring comprising seven BrxC^Aci^ subunits. In that speculative protein model, as well as in the particles observed on negative stain EM images, a pore is present with an inner diameter sufficient to potentially accommodate a DNA duplex (**Figure 3c**). These observations are consistent with our prior observation of higher order, nucleic-acid associated protein complexes in SEC analyses (**Figure 1c**) and the ability of dimeric BrxC^Aci^ to form high molecular weight DNA-bound complexes in EMSA analyses (**Figures 1d, g and h**).

Negative-stain TEM analysis of purified BrxC^Eferg^ also displayed the presence of similar heptameric ring-shaped particles, again at low frequency within a heterogeneous background of protein species (**Supplementary Figure S7**).

### The BrxC N-terminal AAA+ domain also binds BrxB:PglZ

We previously observed that co-expression of *Acinetobacter* and *Salmonella* BrxB and PglZ resulted in stable binary B:Z complexes (19). To determine whether larger *Acinetobacter* BREX assemblies could be detected, we carried out pull-down experiments using purified proteins or various co-expressed combinations of BREX constructs, with only a single construct affinity tagged in each experiment.

In our initial pull-down experiments, we observed the association of BrxB^Aci^, PglZ^Aci^, BrxC^Aci^(E269Q) and PglX^Aci^ (where PglZ^Aci^ contained a twin-strep affinity tag) when all four factors were co-expressed in *E. coli* (**Supplemental Figure S8**), consistent with previous observations in other BREX systems (21,22). The identities of the co-eluting species in that experiment, including PglX^Aci^ and BrxC^Aci^ (which co-migrate by SDS-PAGE) were confirmed by mass spectroscopy. However, this complex did not remain stable on SEC, and we were unable to further structurally characterize the *Acinetobacter* B:C:X:Z complex.

Subsequent co-expression experiments revealed a particularly robust interaction between BrxC^Aci^’s AAA+ N-terminal domain (BrxC^Aci^_1-551_), BrxB^Aci^ and PglZ^Aci^. Co-expression of all three constructs in *E. coli*, with PglZ^Aci^ carrying a Twin Strep affinity tag, followed by affinity purification, proteolytic tag removal and SEC, resulted in co-elution of a stable complex containing all three proteins (**Figure 4a**).

**Figure 4.**
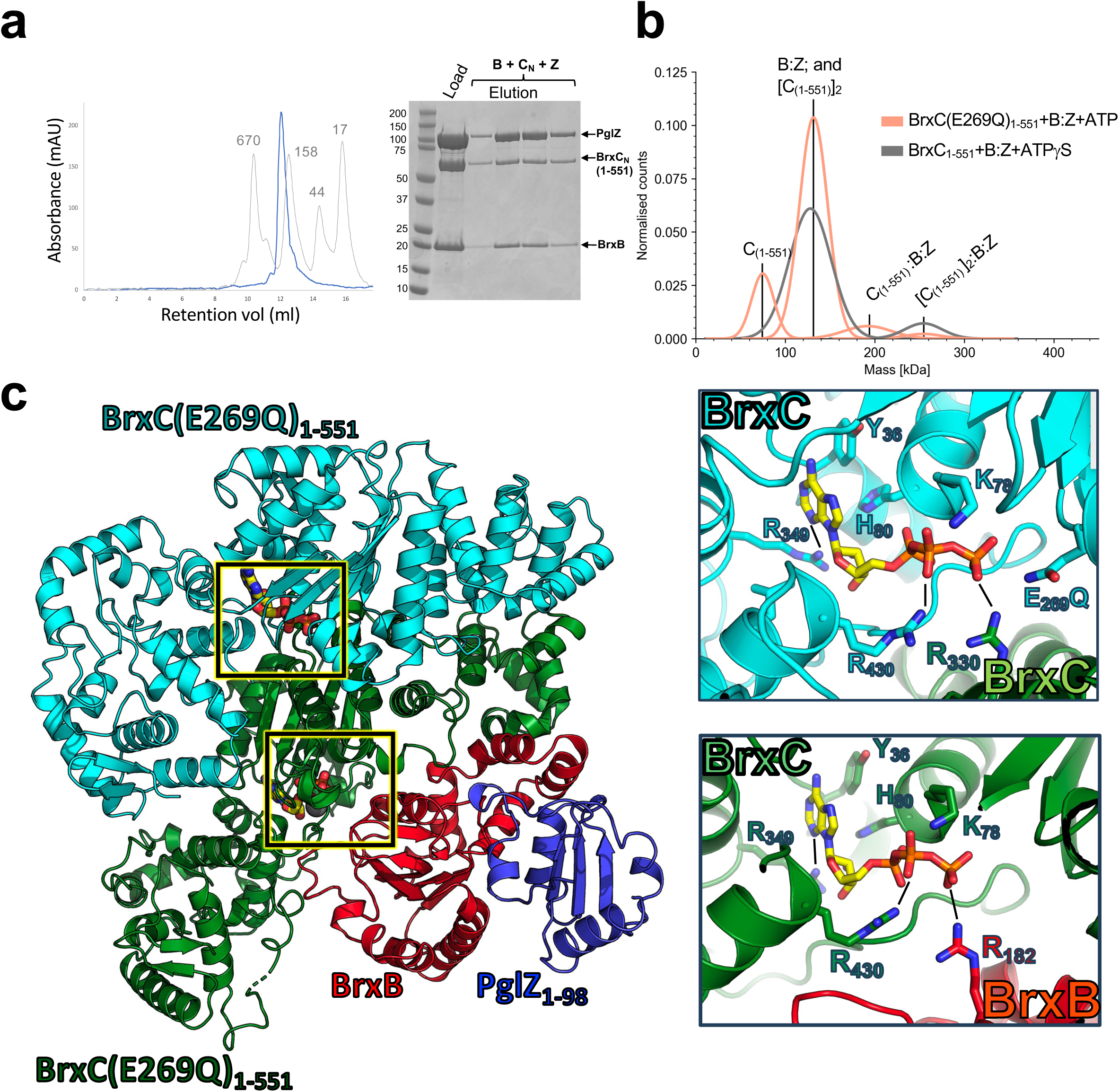
The N-terminal AAA+ domain of *Acinetobacter* BrxC binds the BrxB:PglZ subcomplex. ***Panel a:*** Co-expressed BrxB^Aci^, BrxC^Aci^_1-551_ and strep-tagged PglZ^Aci^ were affinity purified by streptactin chromatography followed by SEC (left trace). All three proteins remain associated through this purification scheme (SDS PAGE, right). ***Panel b:*** Mass photometry analyses of the purified BrxB^Aci^:PglZ^Aci^ dimeric complex (‘B:Z’) incubated with either BrxC^Aci^_1-551_ and ATPγS (gray) or BrxC^Aci^ (E269Q)_1-551_ and ATP (orange). The predicted mass of the B:Z complex is 122 kDa and that of the C_1-551_ monomer is 66 kDa. Complexes consistent with B:Z:(C_1-551_)_1_ and B:Z:(C_1-551_)_2_ stoichiometries are observed. ***Panel c:*** X-ray crystallographic structure of a complex consisting of dimeric BrxC^Aci^(E269Q)_1-551_, full-length BrxB^Aci^ and PglZ^Aci^_1-98_. The BrxC-BrxC interface and the BrxB-BrxC interface (boxes and insets) involve comparable residues that bind a single copy of ATP between the corresponding protein subunits.

Mass photometry analyses using purified proteins confirmed the ATP-dependent formation of complexes with stoichiometries (BrxC^Aci^_1-551_)**_1_**:BrxB^Aci^:PglZ^Aci^ and (BrxC^Aci^_1-551_)_2_:BrxB^Aci^:PglZ^Aci^ (**Figure 4b**), with dimerization of BrxC^Aci^_1-551_ promoted by ATPγS. We next determined a high-resolution structure of a stable ternary complex containing the interacting regions of BrxB^Aci^, BrxC^Aci^ and PglZ^Aci^. In our earlier cryo-EM work attempting to visualize B:Z complexes we were unable to obtain high-resolution reconstructions, likely due to PglZ’s high flexibility from two predicted hinge regions and additional challenges resulting from the preferred orientation of the particles during EM analyses. To mitigate this issue, we co-expressed only PglZ^Aci^’s N-terminal domain (residues 1–98, which mediates its interaction with BrxB^Aci^ (23)), together with BrxC^Aci^(E269Q)_1-551_ and full-length BrxB^Aci^, and isolated a stable complex containing all three polypeptides using the two-step purification strategy described above. The protein complex was crystallized, and its structure was solved using X-ray crystallography at a resolution of ∼2.7 Å (**Supplementary Table S3** and **Figure 4c**). The resulting structure consists of a dimer of BrxC^Aci^(E269Q)_1-551_ domains bound to a single copy of the BrxB^Aci^:PglZ^Aci^_1-98_ heterodimer (corresponding to a ‘(C_1-551_)_2_:B:Z_1-98_’ stoichiometry).

In this complex, two bound ATP molecules and associated divalent cations are observed: one in the same position (**Figure 3a**) visualized previously between the two BrxC^Aci^ subunits (**Figure 4c, top inset)**, and a second in an equivalent position between one subunit of BrxC^Aci^ and BrxB^Aci^ (**Figure 4c, bottom inset**). Whereas in the ATP-binding pocket formed between two BrxC^Aci^ subunits a single arginine residue (R330) is contributed by one subunit, in the analogous pocket between BrxB^Aci^ and BrxC^Aci^, the functionally equivalent residue (R182) is instead contributed by BrxB^Aci^. A key insight from this structural result is that the presence of a bound BrxB^Aci^ subunit would preclude the binding of additional BrxC^Aci^ subunits.

We then repeated the co-expression and purification scheme described above using full-length BrxC^Aci^ (rather than BrxC^Aci^_1-551_) along with BrxB^Aci^ and PglZ^Aci^ (**Figure 5**). Similar to our prior result showing formation of stable complexes containing BrxC^Aci^_1-551_, BrxB^Aci^ and PglZ^Aci^, co-expression of BrxC^Aci^, BrxB^Aci^ and strep-tagged PglZ^Aci^ resulted in co-elution of the three proteins from a streptactin affinity column (**Figure 5a, left**). However, the resulting complex was less stable when purified by SEC, with BrxC^Aci^ largely dissociating from and eluting earlier than the tightly associated BrxB^Aci^:PglZ^Aci^ complex (**Figure 5a, right**).

**Figure 5:**
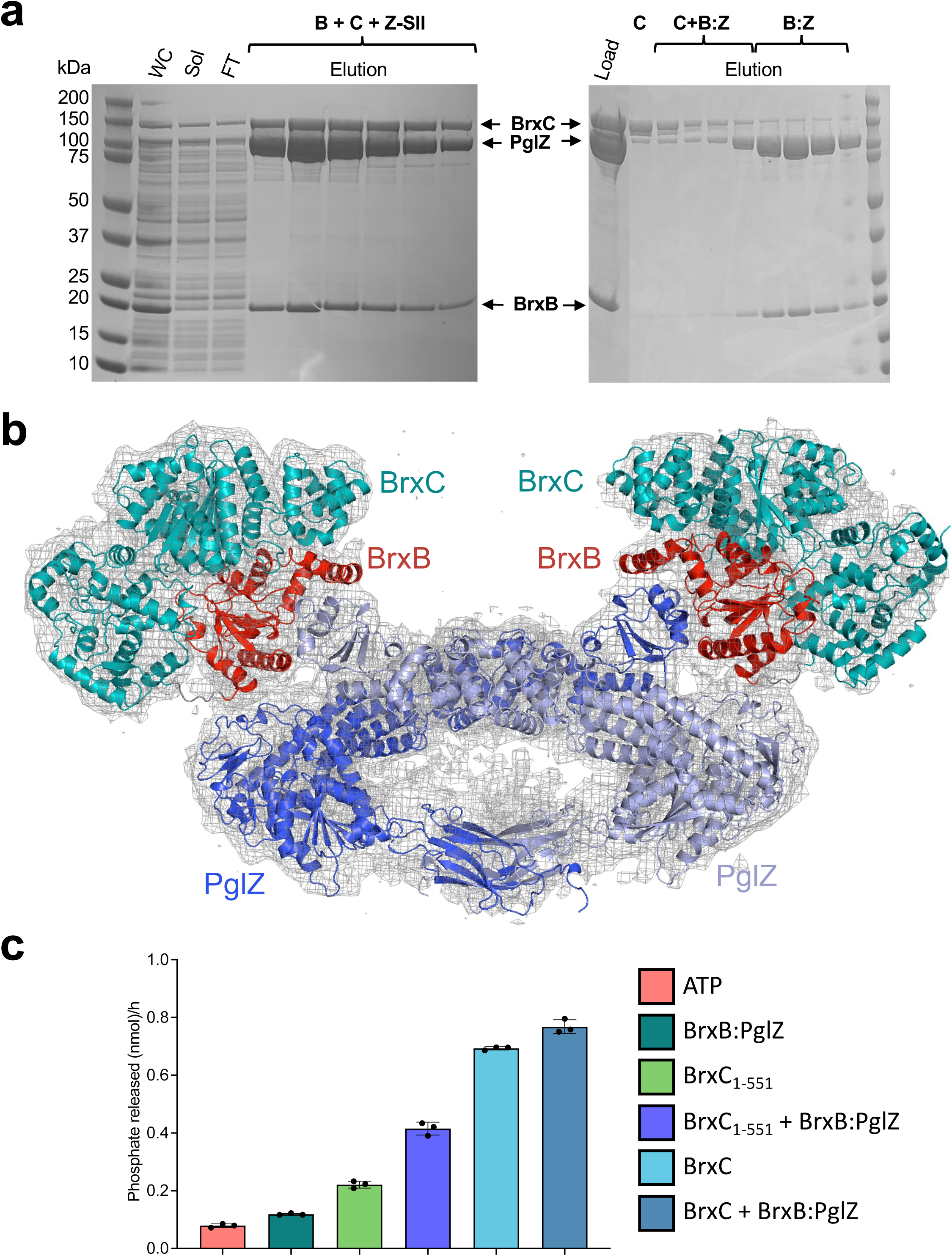
Formation of higher order stoichiometric complexes between full-length *Acinetobacter* BrxC and the BrxB:PglZ subcomplex. ***Panel a, left:*** Co-expression of BrxB^Aci^, strep-tagged PglZ^Aci^ and full-length BrxC^Aci^ demonstrates co-elution of all three subunits by streptactin affinity purification (WC: Whole cell lysate; Sol: filtered soluble lysate; FT: affinity column flow-through). ***Panel a, right:*** The streptacin-eluted B:C:Z complex largely dissociates on SEC into BrxC^Aci^ dimers and the B^Aci^:Z^Aci^ complex. ***Panel b:*** Low resolution cryo-EM analysis of particles corresponding to a complex between a BrxC^Aci^_1-551_-BrxB^Aci^ fusion (teal and red cartoon elements, respectively) and a separate full-length PglZ^Aci^ subunit (blue cartoon) produces a density envelope that agrees closely with a generated predictive model of a BrxC_1-551_-BrxB:PglZ dimer, with the complex held together via interactions between separate ends of each PglZ subunit. ***Panel c:*** ATPase assays of BrxB^Aci^:PglZ^Aci^ and indicated BrxC^Aci^ constructs, comparing each protein alone, or incubated together at 1 µM concentration.

The high-resolution crystal structure of the *Acinetobacter* BrxC_1-551_:BrxB:PglZ_1-98_ ternary complex revealed close proximity between the C-terminus of BrxC^Aci^ and the N-terminus of BrxB^Aci^ (**Figure 4c**). Guided by this observation, we generated a single-polypeptide fusion of BrxC^Aci^_1-553_ and BrxB^Aci^ (‘C-B_fusion_’) and co-expressed it with Strep-tagged full-length PglZ^Aci^. The resulting complex (C-B_fusion_:Z) was purified via affinity chromatography and SEC and was found to remain stable through the final purification step and was therefore subjected to cryo-EM examination.

We observed two unique assemblages in this analysis. The first corresponded to a low resolution density envelope that accommodates our previously established model of the BrxB:PglZ complex (23), along with additional volume adjacent to the BrxB^Aci^ subunit that is consistent with the size of BrxC^Aci^’s N-terminal domain (**Supplementary Figure S1c and d).** The second observed assemblage corresponded to a low resolution (∼7 Å) density map generated from roughly 200,000 particles **(Figure 5b; Supplementary Figure S1e)**, which represented roughly 1/3 of all particles visible within individual micrographs. Two copies of an AlphaFold3–predicted C-B_fusion_:Z model (single-particle confidence: pTM = 0.54, ipTM = 0.7) were docked into the density and individual domains adjusted to fit, where they accounted well for the map features and supported a dimeric assembly. In that model, dimerization appears to be primarily mediated by two regions of PglZ: (1) interactions between the central regions of each PglZ partner, and (2) interactions between PglZ’s Ig-like C-terminal domains. This architecture may provide insight into the organization of higher-order (C_1-551_)_2_:B:Z complexes observed by mass photometry (**Figure 4b)**, although it remains unclear whether BrxC’s C-terminal domains dimerize in the full-length complex or perhaps exist as a mix of dimerized and un-dimerized states.

To investigate if the interactions between BrxC^Aci^, BrxB^Aci^, and PglZ^Aci^ described above are generalizable across homologous BREX systems, we further examined whether similar complexes form between individually purified protein constructs from *E. fergusonii* (**Supplementary Figure S9**). Both BrxC^Eferg^_1-551_ and BrxC^Eferg^(E268Q)_1-551_ incubated with a mixture of BrxB^Eferg^ and PglZ^Eferg^ produced two early eluting peaks by SEC (**Supplementary Figure S9a and c**). Analysis of the early peak fractions by SDS-PAGE shows that in the absence of ATP, BrxC_1-551_ elutes at low levels alongside BrxB^Eferg^ and PglZ^Eferg^ (**Supplementary Figure S9b**), and the presence of ATP and MgCl_2_ abolishes this interaction (**Supplementary Figure S9b, bottom**). Comparatively higher amounts of BrxC^Eferg^(E268Q)_1-551_ were observed alongside BrxB^Eferg^ and PglZ^Eferg^ within the early eluting peak fractions both in the absence (**Supplementary Figure S9d, top**) and presence (**Supplementary Figure S9d, bottom**) of ATP and MgCl_2_. These data further support a model where (i) the AAA+ domain of BrxC forms a complex with BrxB and PglZ, (ii) the interaction is dependent upon or stabilized by the binding, but not hydrolysis, of ATP, and (iii) the interaction is conserved between homologous BREX systems found in different bacteria.

Having identified an ATP-dependent interaction between BrxC’s AAA+ domain, BrxB, and PglZ, we sought to analyze the effect, if any, that the interaction has on the ATPase activity of BrxC (**Figure 5c and Supplementary Figure S10**). We first tested the effect of BrxB^Aci^:PglZ^Aci^ on full-length BrxC^Aci^ and observed little to no change in ATPase activity. Because ATPase activity in the full-length protein may be influenced by dimerization of BrxC’s AAA+ domains, we next tested the BrxC_1-551_ constructs from both species to better isolate the contribution of the B:Z complex. Under these conditions the B:Z complex induced a ∼2-6 fold increase in ATPase activity (**Figure 5c and Supplementary Figure S10**).

To determine the individual contributions of BrxB and PglZ to the observed increase in BrxC_1-551_ ATPase activity, we tested both *E. fergusonii* factors alone (**Supplementary Figure S10**; BrxB^Aci^ could not be tested because it is only stable when co-purified with PglZ^Aci^). Incubation with BrxB^Eferg^ alone induced a ∼4-fold increase in ATPase activity of BrxC^Eferg^_1-551_ compared to each individual component, whereas the observed activity of BrxC^Eferg^_1-551_ incubated with PglZ^Eferg^ was comparable to the additive contribution of the individual components rather than an increase. This effect was further amplified by the BrxB^Eferg^:PglZ^Eferg^ complex, reaching a ∼6-fold increase in ATPase activity. BrxB^Eferg^ and BrxB^Eferg^:PglZ^Eferg^ also induced a small increase in the ATPase activity of BrxC^Eferg^(E268Q)_1-551_. These results indicate that the B:Z complex can regulate BrxC ATPase activity.

### Characterization of mutants that individually disrupt BrxC:BrxC or BrxB:BrxC association

Given that BrxB associates with BrxC through the same interface that mediates higher-order BrxC multimerization—suggesting that these two binding events are mutually exclusive—we identified a mutation in each subunit designed to selectively disrupt one interaction without affecting the other. In the BrxB^Aci^-BrxC^Aci^ (“B-C”) interface, BrxB^Aci^ residue R82 forms a well-ordered electrostatic contact to D204 in BrxC^Aci^ (**Figure 6a**). Conversely, in the BrxC^Aci^-BrxC^Aci^ (“C-C”) interface, R330 in one subunit interacts with bound ATP and with E289 in the other subunit but is not involved in the B-C interface.

**Figure 6:**
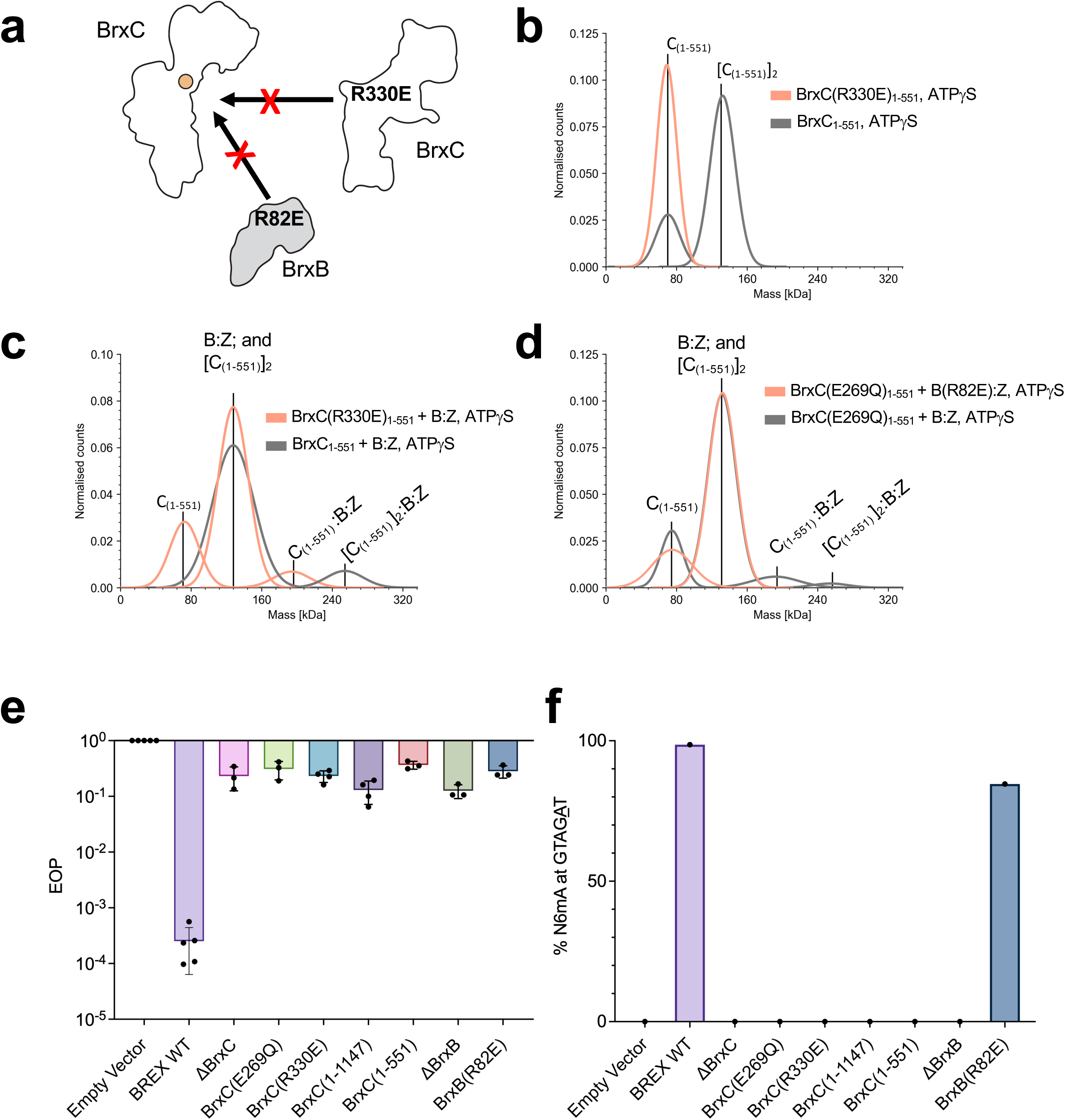
Disruption of BrxC:BrxC and BrxB:BrxC interfaces, and impact on *Acinetobacter* BREX activity. ***Panel a:*** Schematic depicting the BrxC^Aci^(R330E) and BrxB^Aci^(R82E) mutants predicted to disrupt the BrxC:BrxC and BrxB:BrxC interfaces, respectively. ***Panel b:*** Mass photometry analysis of BrxC^Aci^_1-551_ (gray) and BrxC^Aci^(R330E)_1-551_ (orange) in the presence of ATPγS shows that the R330E mutant blocks BrxC^Aci^ N-terminal self-association. ***Panel c:*** The B^Aci^:Z^Aci^ complex was incubated with BrxC^Aci^_1-551_ (gray) or BrxC^Aci^(R330E)_1-551_ (orange) in the presence of ATPγS. BrxC^Aci^(R330E)_1-551_ does not disrupt the BrxB-BrxC interaction but only forms a complex with B:Z in a B:Z:(C_1-551_)_1_ stoichiometry (i.e 1:1:1). ***Panel d:*** BrxC^Aci^(E269Q)_1-551_ was incubated with BrxB^Aci^:PglZ^Aci^ or BrxB^Aci^(R82E):PglZ^Aci^ in the presence of ATPγS. This analysis shows that BrxB^Aci^(R82E) abrogates complex formation between B:Z and C_1-551_*. **Panel e:*** Efficiency of Plating (EOP) results for lambda phage challenges against *E. coli* transformed with the indicated *Acinetobacter* BREX constructs and empty vector control. Error bars represent standard deviation from the mean of biological replicates. The truncations of BrxC^Aci^ and mutations disrupting the C-C or B-C interfaces are comparable to deletion of either gene individually and result in nearly complete loss of BREX-mediated protection. ***Panel f:*** PacBio genome methylation data of the same BREX constructs tested by EOP in **panel e**. Sequencing percentage corresponds to the amount of methylated GTAG**<u>A</u>**T N6mA BREX motifs.

We therefore purified variants carrying individual mutations—BrxB^Aci^(R82E) and BrxC^Aci^(R330E)_1-551_—and examined their ability to bind the corresponding WT counterparts using mass photometry. We used the truncated BrxC^Aci^_1-551_ construct for this analysis to uncouple dimerization of the N-terminal AAA+ domain from dimerization mediated by the C-terminal domain present in the full-length protein. As predicted, both charge reversal mutations selectively abrogated their corresponding interactions: BrxC^Aci^(R330E)_1-551_ selectively eliminated C-C self-association (**Figure 6b**) but retained interaction with B^Aci^:Z^Aci^, albeit with only one copy of BrxC^Aci^_1-551_ (forming a BrxC_1-551_:B:Z complex with 1:1:1 stoichiometry) (**Figure 6c**). BrxB^Aci^(R82E) abolished the B-C interaction (**Figure 6d**).

We also tested how these individual mutants affected BrxC^Aci^_1-551_ ATPase activity (**Supplementary Figure S10)**. BrxC^Aci^(R330E)_1-551_ exhibited ATPase activity comparable to BrxC^Aci^_1-551_ alone and showed a similar increase in ATPase activity when incubated with BrxB^Aci^:PglZ^Aci^, consistent with the BrxC(R330E) mutant not affecting the B-C interface. In contrast, BrxB^Aci^(R82E):PglZ^Aci^ failed to stimulate BrxC^Aci^(WT)_1-551_ ATPase activity, consistent with this mutant disrupting the B-C interface.

### Impact of BREX mutations on phage restriction and methylation

Finally, we measured the impact of our studied mutations on the two biological outcomes of BREX activity, (i) phage restriction, and (ii) methylation of the genomic PglX^Aci^ target site (GTAG**<u>A</u>**T) when expressed in *E. coli*. Whereas BREX^Aci^ WT conferred 3-4 orders of protection against lambda phage challenge relative to an empty vector control, all mutants lost the ability to restrict lambda (**Figure 6e**). In the corresponding methylation analysis, BREX^Aci^ WT conferred nearly complete methylation of PglX^Aci^ sites in the *E. coli* ER2796 (35) genome (98.6%) (**Figure 6f**). All constructs except the BrxB^Aci^(R82E) mutant demonstrated complete loss of methylation activity; the BrxB (R82E) mutant conferred 84.6% coverage (**Figure 6f**). Although slightly reduced relative to BREX^Aci^ WT, BrxB^Aci^(R82E) mutant displays an uncoupling of phage protection and methylation previously observed for point mutations in *Salmonella* BrxC’s C-terminal domain (37).

Corresponding analysis with BREX^Eferg^ produced a different result; instead of being inactivated, phage restriction was still observed for BrxC^Eferg^(E268Q) and an intermediate restriction phenotype was observed for BrxC^Eferg^_1-551_ (**Supplementary Figure S11a**). It was noted that transformation efficiency to produce both these mutant strains was very low, and the cells grew slowly. Interestingly, methylation analysis of both mutations showed no BREX methylation activity (**Supplementary Figure S11b**). Lack of methylation would likely account for the slow growth of these strains if BREX remained sufficiently active to protect against phages while also being modestly toxic to the unmethylated host. This result reflects another nuance in BREX activity between homologous systems.

## DISCUSSION

We and others favor a model in which overlapping subsets of BREX proteins assemble into distinct complexes responsible for either host chromosome methylation or phage restriction (18,21,26,38), with regulatory mechanisms coordinating dynamic transitions between these functional states. Here we observe that the BREX^Aci^ proteins BrxB, BrxC, PglZ and PglX form a complex (“B:C:Z:X”) when co-expressed in *E. coli* (**Supplementary Figure S8**); analogous complexes have also been reported in the *E. coli* and *Salmonella* type 1 BREX systems (21,22). Given the recently demonstrated nuclease activity of PglZ (23), we envision that phage infection induces remodeling of a methylation-specific complex into a restriction complex, thereby activating PglZ nuclease activity and potentially other BREX factors. The roles of BrxA and BrxL, which are required for restriction in some systems but dispensable in others (18,26,27) remains unresolved. Our study highlights the formation of B:C:Z complexes that may serve as a “base” for B:C:Z:X assemblies; and a potential central role for BrxC in (1) potentiating methylation and restriction and (2) serving as a key point of regulation controlling the transition between these activities. The various structural states described in this study can be summarized (**Figure 7**).

**Figure 7.**
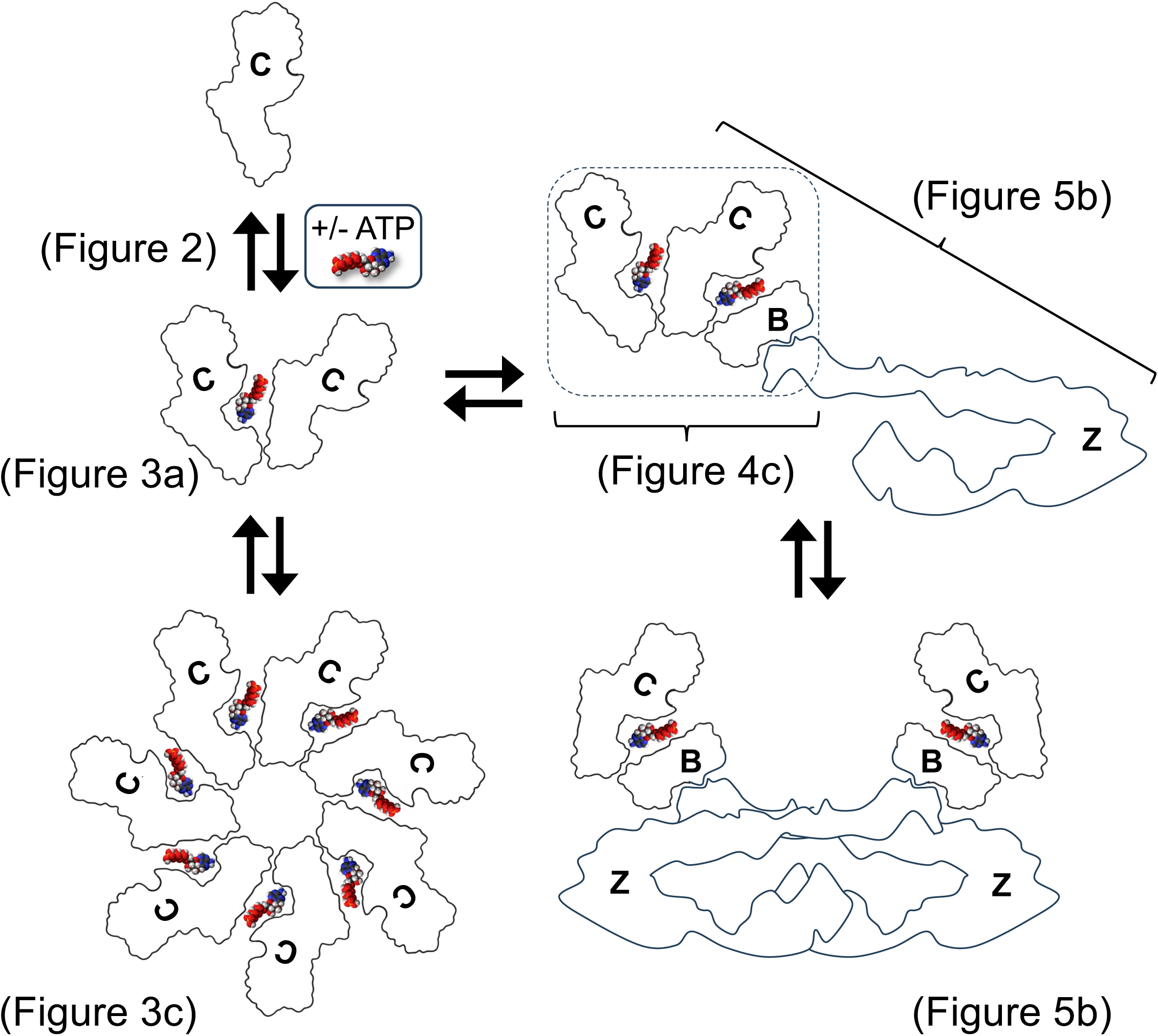
BrxC forms multiple assemblies. The schematic depicts the various structures formed by BrxC, based on findings in the indicated figures. For clarity, only BrxC’s N-terminal domain is represented.

BrxC can self-assemble into at least two distinct structural states (**Figures 3a,c and 7**). First, it forms a homodimer anchored by a tight C-terminal dimerization domain that creates a stable hinge, while its N-terminal AAA+ ATPase domains undergo reversible, ATP-regulated dimerization in an asymmetric “spooning” configuration, with key contacts formed between residues from one BrxC subunit and the ATP-binding pocket of its partner (**Figure 3a**). A second structural state consists of higher order heptameric assemblies—observed with BrxC homologs from two different species—containing a central pore compatible with DNA binding (**Figure 3c**). AF3 models consistent with these structures predict that the same interface that mediates BrxC AAA+ dimerization could be used by additional subunits to generate higher order assemblies, including heptamers that we observe in cryo-EM analyses.

We further demonstrate that BrxC associates with the stable PglZ:BrxB dimer through interactions centered on BrxC’s ATPase domain and BrxB’s inactivated AAA+ domain and can assemble into B:Z:C_2_ or (B:Z)_2_:C_2_ stoichiometries (**Figures 4, 5 and 7**). In these configurations, BrxC forms a stable dimer mediated by its C-terminal domains, and its N-terminal domains are accessible for binding to BrxB. We further visualized a low resolution (C-B_fusion_)₂:Z₂ hybrid complex lacking BrxC’s C-terminal domain that indicates two points of self-dimerization for PglZ (**Figure 5b)**. Combined, these results show that B:C:Z form a core complex that may assume various stoichiometries and suggest two structural roles for BrxB: (1) regulating higher-order BrxC oligomerization through its interaction with BrxC’s AAA+ domain, and (2) bridging BrxC and PglZ within B:C:Z complexes.

In addition to demonstrating the formation of B:C:Z assemblages, we show that BrxB:PglZ regulates two activities of BrxC. First, BrxB:PglZ increases the ATPase activity of BrxC’s N-terminal ATPase domain, as shown when incubated with a BrxC_1-551_ construct that uncouples BrxC dimerization from ATPase activity (**Figure 5c**). BrxB’s boost to BrxC’s ATPase activity may perhaps be due to facilitating ADP release, since BrxB^Eferg^ increases both BrxC^Eferg^_1-551_ and BrxC^Eferg^(E268Q)_1-551_ activity (**Supplementary Figure S10**). Second, BrxB:PglZ augments BrxC association with DNA (**Figure 1h**). We also show that BrxA both binds BrxC and inhibits BrxC association with DNA (**Figure 1h**). These findings indicate that BrxA and BrxB—recently shown to be universally present across all BREX subtypes (17)—may play important regulatory roles through their interactions with BrxC.

Genetic data indicate that the BrxB:BrxC:PglZ:PglX (B:C:Z:X) module is required for both the restriction and modification functions of the BREX system(18,26,27). Consistent with this, mutations in the M.StyLT7II_BREX system have been identified within the C-terminal regions of PglX (39) and BrxC (37) that uncouple BREX-mediated methylation and restriction activities, likely by disrupting interactions with other BREX factors. Here, we show that a BrxB^Aci^(R82E) mutant that disrupts the BrxC-BrxB interface—demonstrated *in vitro* by both loss of binding and failure to augment BrxC^Aci^_1–551_ ATPase activity—also uncouples these activities, remaining largely competent for methylation but deficient in restriction. Together, these results provide functional evidence supporting a role for BrxB in regulating the coordination between BREX methylation and restriction activities through its interaction with BrxC.

Certain functional domains and enzymatic activities are repeatedly found across a wide range of bacterial defense systems, in accordance with the “guns for hire” paradigm in which enzymes have shuttled between mobile genetic elements (MGEs) and defense systems throughout microbial evolution (40,41). BREX systems have a high diversity of protein domains and activities, but are also notable for repeatedly using certain domains—for example the iSTAND and wHTH domains (17). A recent bioinformatic analysis identified a family of BrxC homologs (DUF499) that retain strong N-terminal homology with canonical BREX BrxC proteins (DUF6079) but differ in their C-terminal regions (17). The DUF499 BrxC variants co-occur with a PglX methyltransferase homolog and a helicase, and the locus invariably encodes an associated endonuclease. This three-gene configuration likely constitutes a bacterial defense system and has been termed a BREX-related (“BR”) system (17). Of the three recognized BR subtypes, the type 3 subsystem lacks an endonuclease and instead encodes a minimal PglZ phosphoesterase domain together with a BrxB homolog—strongly supporting a functional role for PglZ’s DNA cleavage activity and the B:Z modules in BREX systems. BrxC homologs are also present in Dnd and Ssp systems (17,42), highlighting BrxC as a conserved component across multiple bacterial defense systems. This prevalence is further highlighted by the phage-encoded BREX inhibitor OrbA that has been shown to inhibit BREX function by specifically targeting BrxC (43), which provides evidence for a central role of BrxC in mediating defense activity.

Our presented structures and functional data into the activity of conserved BrxC proteins provide another level of detail to the complex and elusive BREX mechanism. The increased identification of BREX-like functions across multiple defenses and the generalized nature of our findings will provide future opportunities to develop a fuller overview of BREX activity.

## DATA AVAILABILITY

The crystal structure of BrxC^Aci^_1-551_:BrxB ^Aci^:PglZ ^Aci^_1-98_ has been deposited in the RCSB database (PDB ID code 9ZLL). The following cryo-EM models and corresponding maps have been deposited in the PDB and EMBD databases and will be publicly released April 8: BrxC ^Aci^(E269) dimer (PDB code 9ZDX / EMD-74076), BrxC^Aci^_1-551_-BrxB^Aci^ fusion:PglZ^Aci^ dimer (PDB code 9ZN5 / EMD-74435), BrxC^Aci^_1-551_-BrxB^Aci^ fusion:PglZ^Aci^ monomer (EMD-74400).

## ACKNOWLEDGEMENTS

This research was supported by the Electron Microscopy Shared Resource, RRID:SCR_022611, of the Fred Hutch/University of Washington/Seattle Children’s Cancer Consortium (P30 CA015704). Cryo-EM molecular graphics and analyses were performed with UCSF ChimeraX, developed by the Resource for Biocomputing, Visualization, and Informatics at the University of California, San Francisco, with support from National Institutes of Health R01-GM129325 and the Office of Cyber Infrastructure and Computational Biology, National Institute of Allergy and Infectious Diseases. We thank Drs. Theo Humphreys, Anvesh Dasari, and Melody Campbell at the FHCC Electron Microscopy Core facility for helpful discussions and advice, and Gerald Smith for providing λ phage for restriction analyses.

The Berkeley Center for Structural Biology is supported in part by the Howard Hughes Medical Institute. The Advanced Light Source is a Department of Energy Office of Science User Facility under Contract No. DE-AC02-05CH11231. The Pilatus detector on 5.0.1. was funded under NIH grant S10OD021832. The ALS-ENABLE beamlines are supported in part by the National Institutes of Health, National Institute of General Medical Sciences, grant P30 GM124169.

## AUTHOR CONTRIBUTIONS

A.J.K, J.J.R. and L.A.D. contributed equally. A.J.K, L.A.D., S.F.M, J.P.A. D.V. and B.K.K produced and analysed *Acinetobacter* proteins. J.J.R. and A.K. expressed *E. fergusonii* proteins and performed biochemical analyses. L.A.D. performed cryo-EM practical aspects, data collection and data processing for *Acinetobacter* proteins. A.J.K. conducted phage assays testing variants of *Acinetobacter* BREX. M.P. conducted phage assays testing variants of *E.ferg* BREX.

*L.A-B and E.A-P* performed cryo-EM practical aspects, data collection and data processing for *E. ferg* BrxC. Y.A.L., T.R.B., M.P. and A.N. carried out PacBio sequencing and analysis. D.L.S., E.A-P, B.L.S., B.K.K., and T.R.B. supervised the project and obtained funding. All authors contributed to data analysis and writing the manuscript

