## Supplemental Figures for "Competing forms of protein-protein association and DNA binding exhibited by BrxC from the BREX phage restriction system"

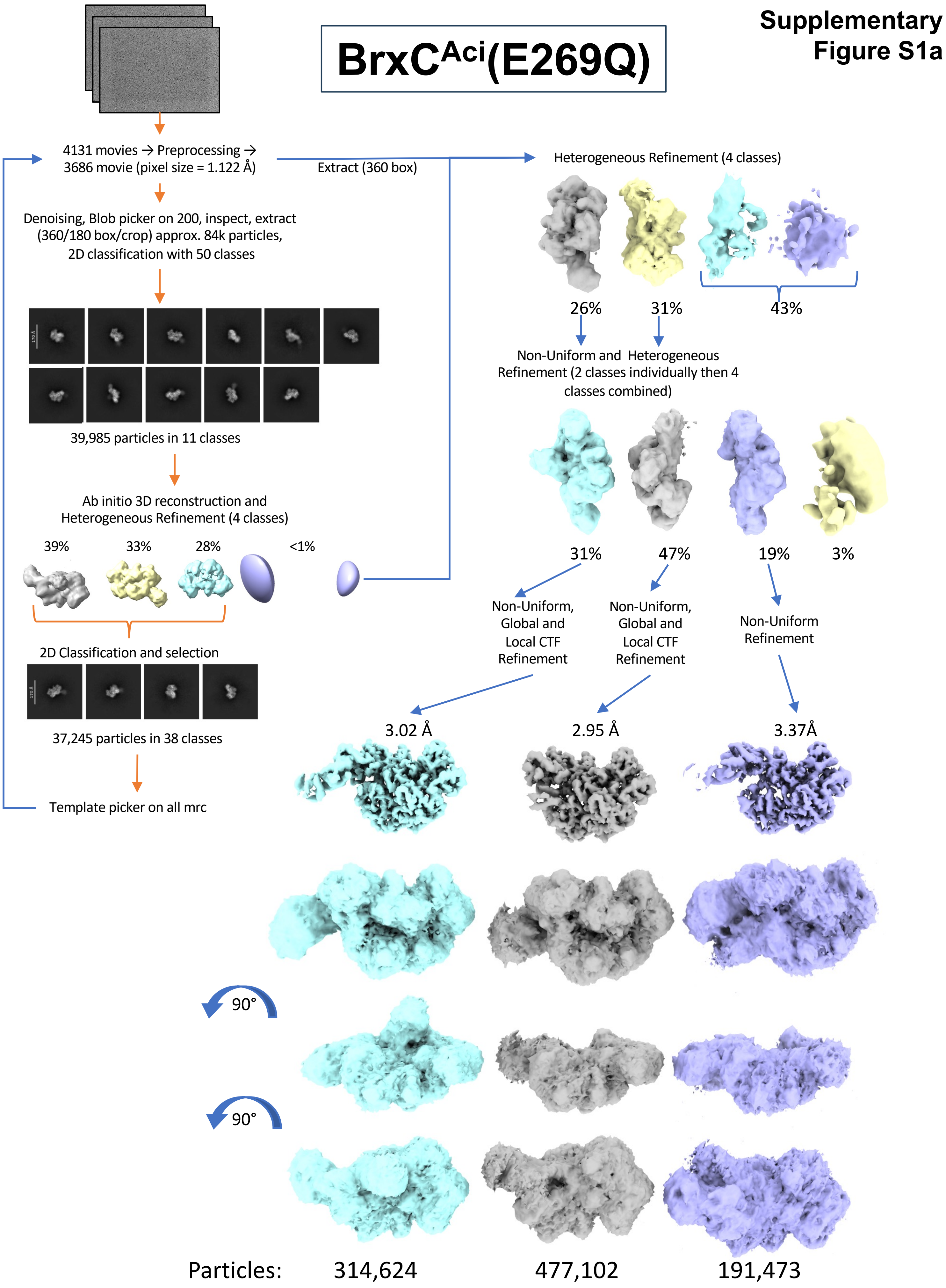
Supplementary Figure S1a
BrxC$^{Aci}$(E269Q)
4131 movies → Preprocessing → 3686 movie (pixel size = 1.122 Å)
Extract (360 box)
Heterogeneous Refinement (4 classes)
26%
31%
43%
Non-Uniform and Heterogeneous Refinement (2 classes individually then 4 classes combined)
31%
47%
19%
3%
Non-Uniform, Global and Local CTF Refinement
Non-Uniform, Global and Local CTF Refinement
Non-Uniform Refinement
3.02 Å
2.95 Å
3.37Å
Denoising, Blob picker on 200, inspect, extract (360/180 box/crop) approx. 84k particles, 2D classification with 50 classes
39,985 particles in 11 classes
Ab initio 3D reconstruction and Heterogeneous Refinement (4 classes)
39%
33%
28%
<1%
2D Classification and selection
37,245 particles in 38 classes
Template picker on all mrc
90°
90°
Particles: 314,624 477,102 191,473

### BrxC$^{Aci}$(E269Q)

**Supplementary Figure S1b**

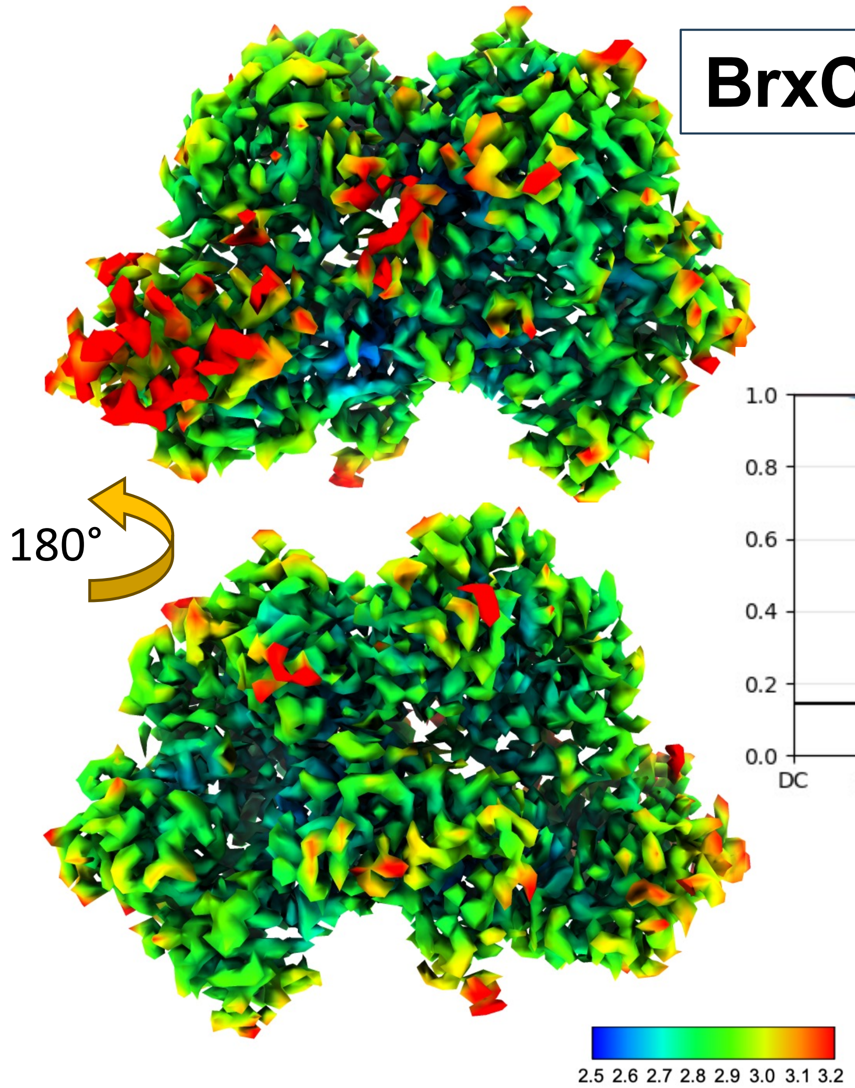


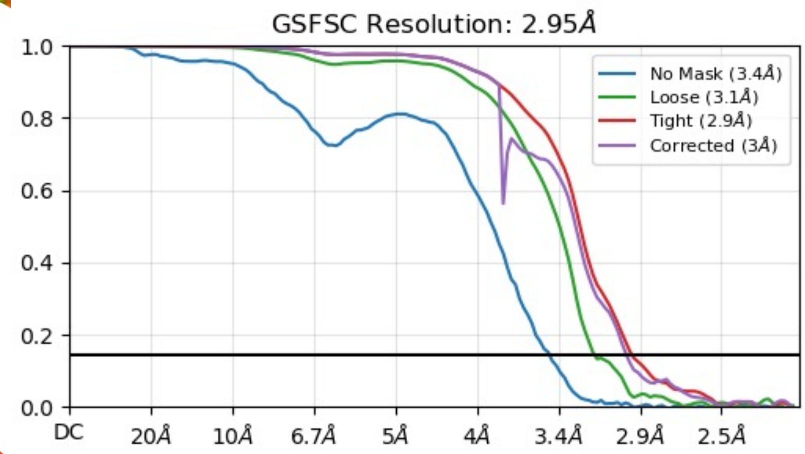


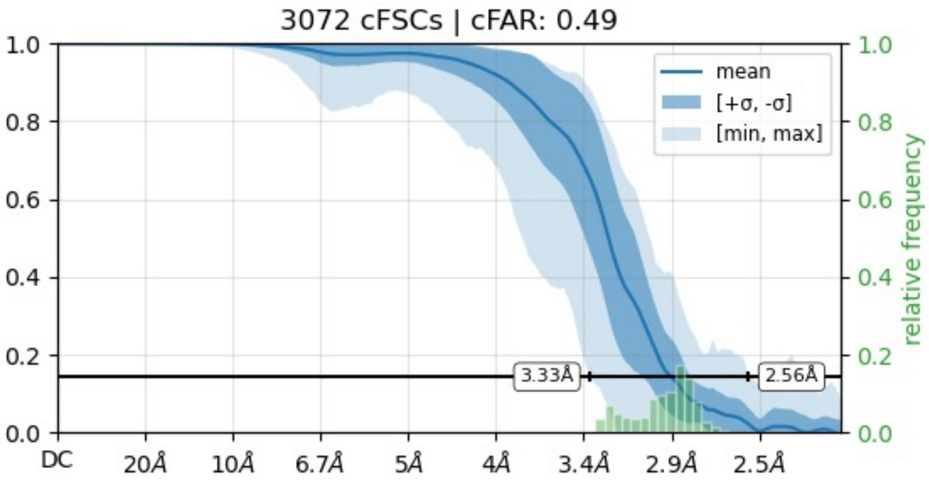


Orientation diagnostics

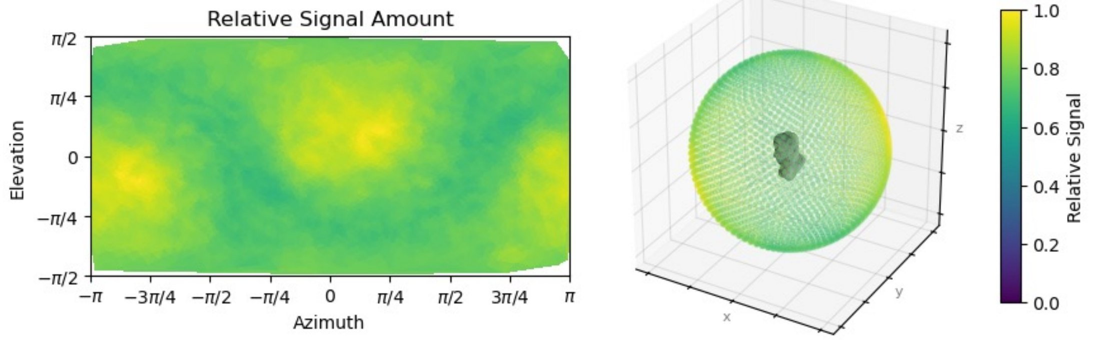

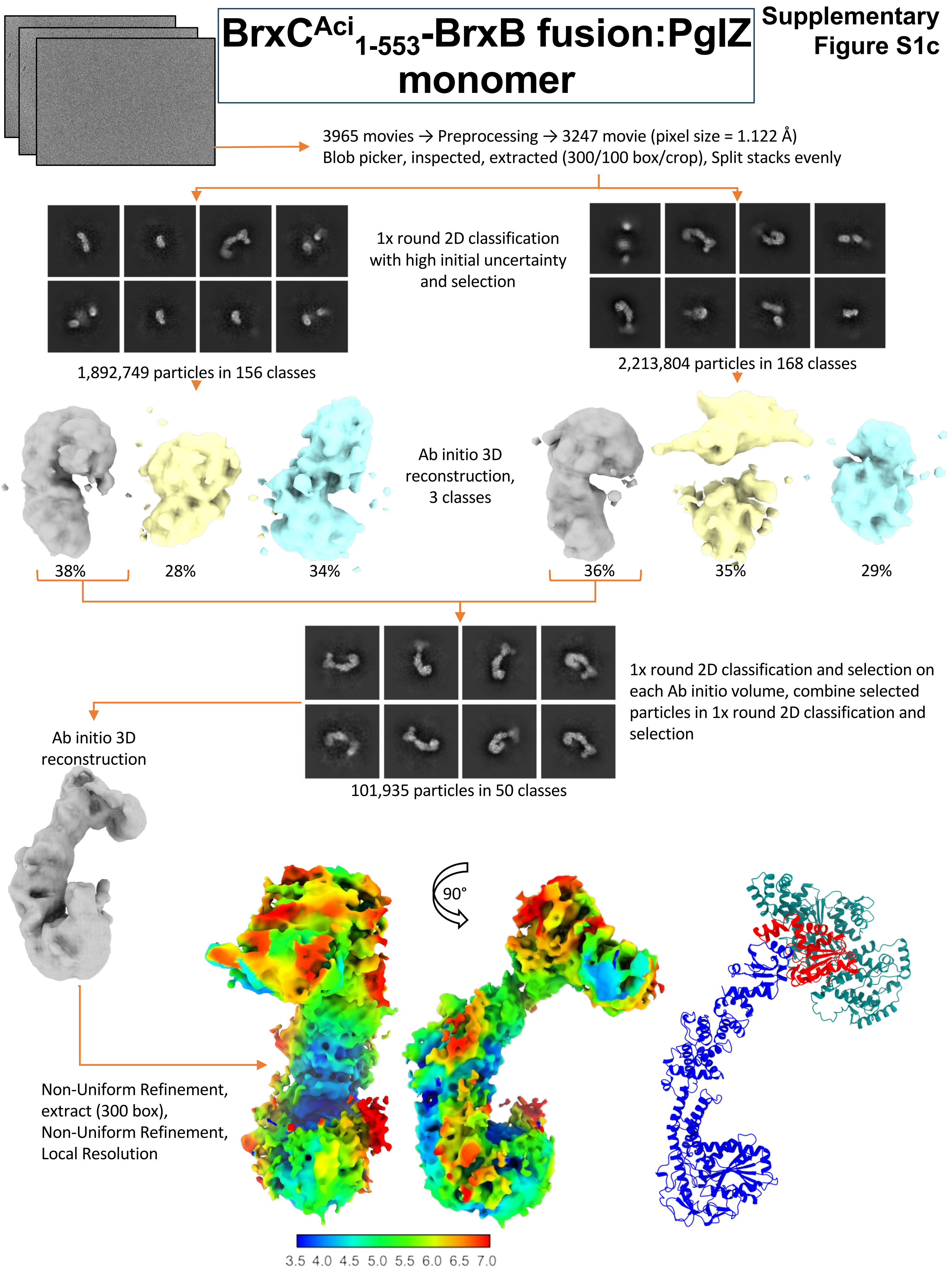

BrxC^Aci_1-553-BrxB fusion:PglZ monomer
Supplementary Figure S1c
3965 movies → Preprocessing → 3247 movie (pixel size = 1.122 Å)
Blob picker, inspected, extracted (300/100 box/crop), Split stacks evenly
1x round 2D classification with high initial uncertainty and selection
1,892,749 particles in 156 classes
2,213,804 particles in 168 classes
Ab initio 3D reconstruction, 3 classes
38%
28%
34%
36%
35%
29%
1x round 2D classification and selection on each Ab initio volume, combine selected particles in 1x round 2D classification and selection
101,935 particles in 50 classes
Ab initio 3D reconstruction
90°
Non-Uniform Refinement, extract (300 box), Non-Uniform Refinement, Local Resolution
3.5 4.0 4.5 5.0 5.5 6.0 6.5 7.0

### BrxC$^{Aci}{}_{1\text{-}553}$-BrxB fusion:PglZ monomer

**Supplementary Figure S1d**

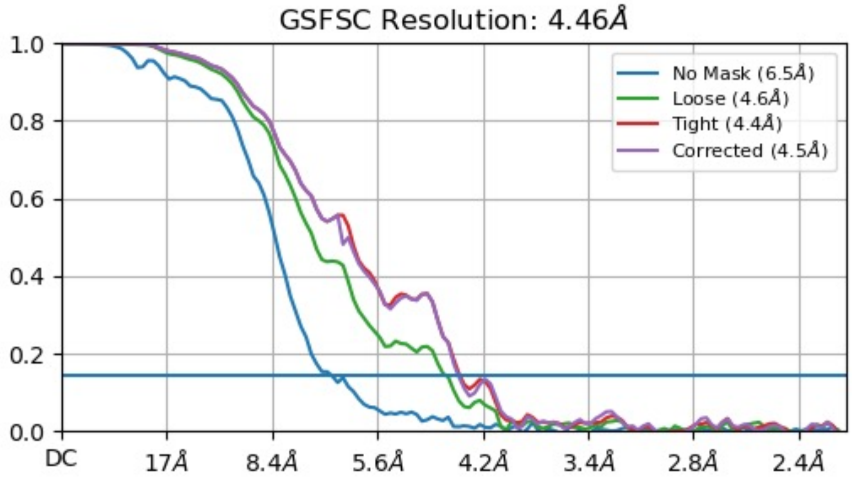


2D Classification examples

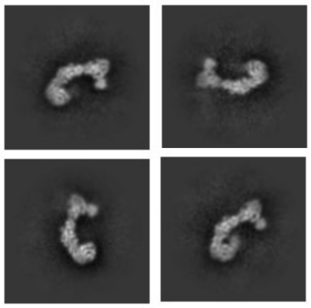

BrxC-BrxB/PglZ complex

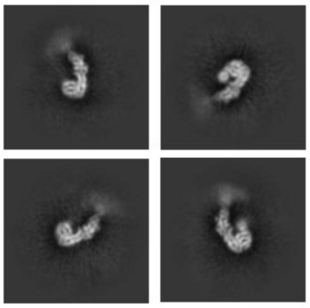

C-terminal PglZ

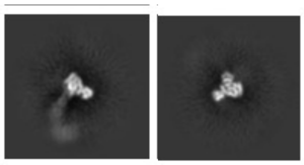

BrxC-BrxB fusion
(~8% of particles)

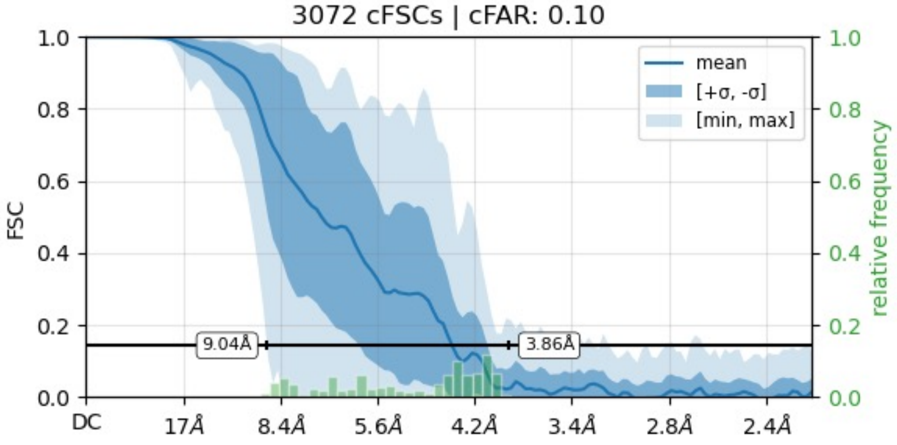


Orientation diagnostics

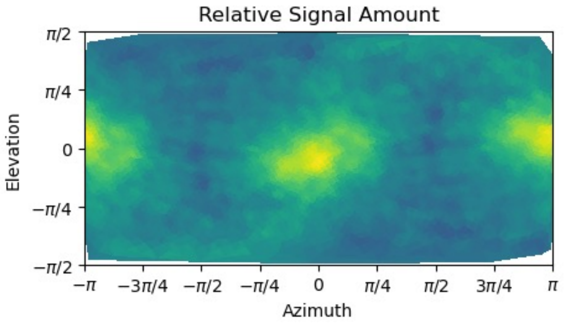


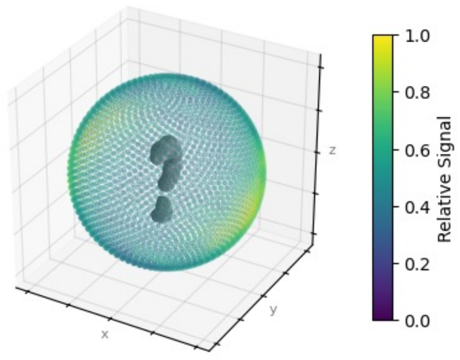

**Supplementary Figure S1e**

### BrxC$^{Aci}_{1\text{-}553}$-BrxB fusion: PglZ dimer

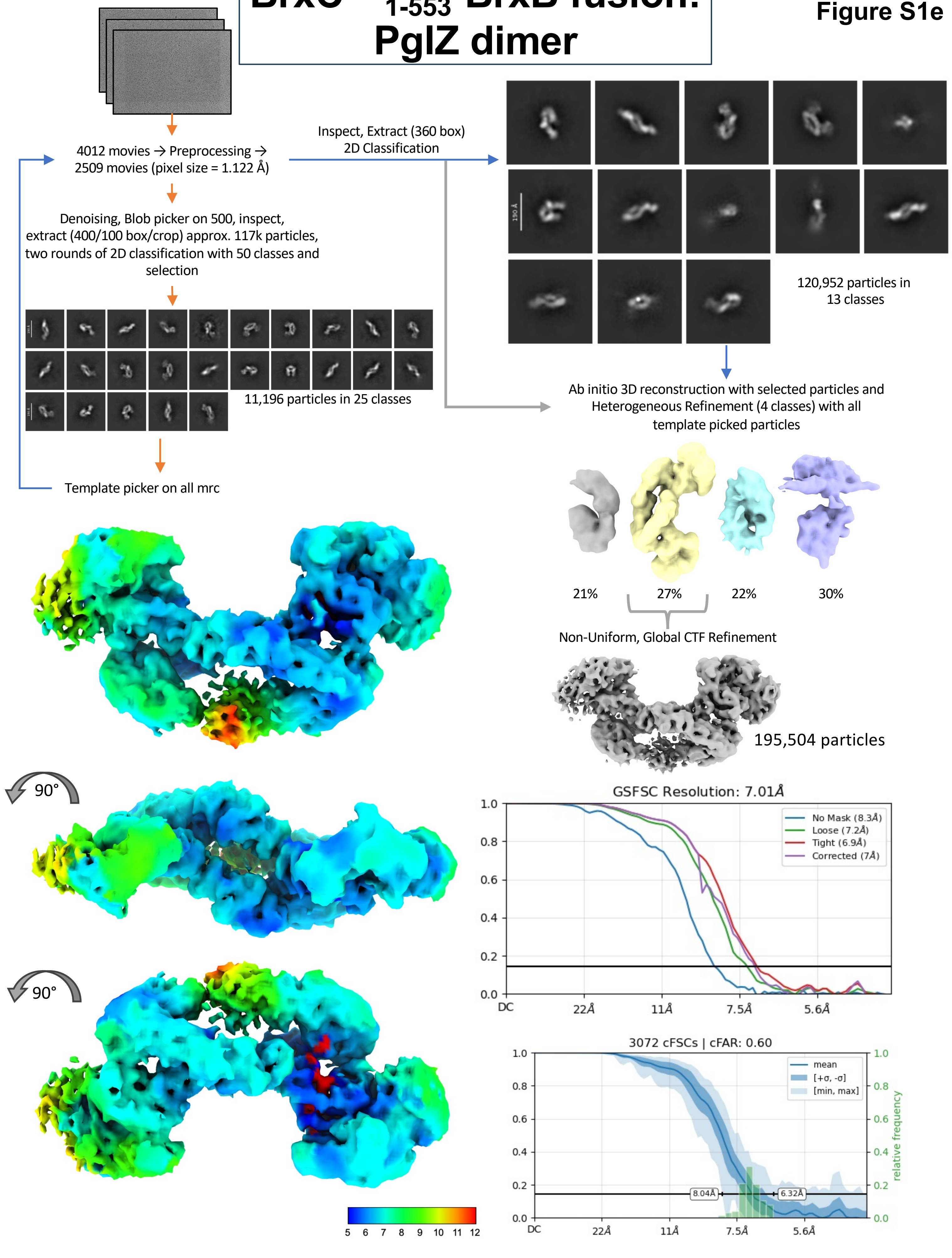

**Supplementary Figure S1**. **Schematic of structural analyses of various *Acinetobacter* BrxC assembly states via cryo-EM**.

***Panel a:*** Data collection and processing flow chart for single particle analyses of the $BrxC^{Aci}$(E269Q) dimer. Example images of a subset of selected 2D Classes are presented showing various orientations of the particles. The three distinct 3D volumes are show at both high and low contour levels to highlight the differences between them.

***Panel b:*** Local resolution, GSFSC resolution, and orientation analysis of final map reconstruction for the $BrxC^{Aci}$(E269Q) dimer.

***Panel c:*** Data collection and processing flow chart for single particle analysis of the $BrxC_{1\text{-}553}$-$B_{fusion}^{Aci}$:$Z^{Aci}$ complex at a 1:1 stoichiometry. Example images of a subset of selected 2D Classes are presented showing various domain alignments and orientations of the particles. Local resolution of the final volumes is shown at the bottom with a ribbon diagram of an AlphaFold3 model roughly fit to the volume using ChimeraX Fit in Map utility. $BrxC_{1\text{-}553}$-$B_{fusion}^{Aci}$ and $PglZ^{Aci}$ residues 1-98 were treated as one domain during the fitting.

***Panel d:*** GSFSC resolution, 2D classification examples, and orientation analysis of final map reconstruction of the $BrxC_{1\text{-}553}$-$B_{fusion}^{Aci}$:$PglZ^{Aci}$ complex at a 1:1 stoichiometry. ***Panel e:*** Data collection and processing flow chart for single particle analyses of the $BrxC_{1\text{-}553}$-$B_{fusion}^{Aci}$:$PglZ^{Aci}$ dimer, with the constructs present in a 2:2 stoichiometry. Local resolution, GSFSC resolution and cFAR are present in the lower half of the panel.

### Supplementary Figure S2

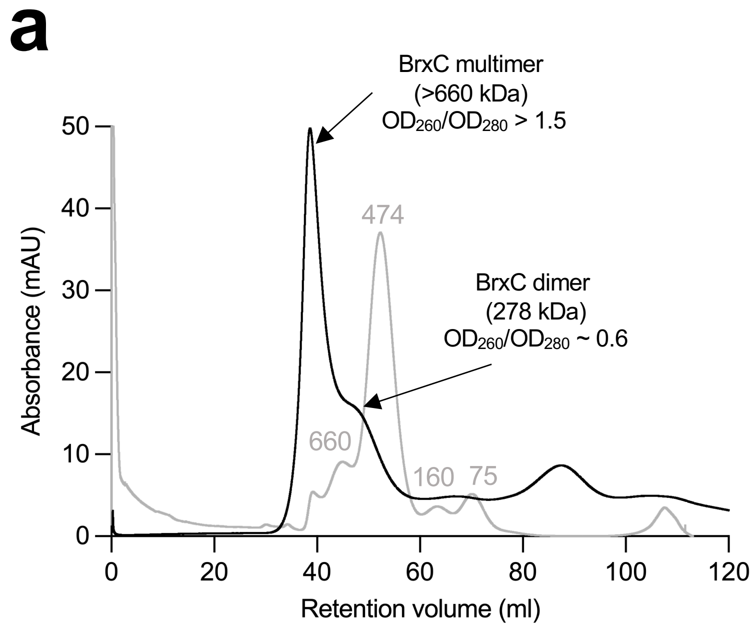


**Supplementary Figure S2. Solution behavior of BrxC from *E. fergusonii*.**
***Panels a and b:*** Size exclusion chromatography using a HiPrep™ 16/60 Sephacryl® S-300 HR column for $BrxC^{Eferg}$ (**a**) and $BrxC^{Eferg}$(E268Q) (**b**). The traces represent the absorbance at 280 nM. The peaks comprising the $BrxC^{Eferg}$ and $BrxC^{Eferg}$(E268Q) multimer and dimer are highlighted by black arrows. The calculated mass (kDa) of the dimer and the expected size of the multimer, based on the size of the largest calibration protein, are shown. The $OD_{260}/OD_{280}$, measured by nanodrop, are also shown. The grey trace represents the calibration using conalbumin (75 kDa), aldolase (160 kDa), ferritin (474 kDa), thyroglobulin (660 kDa), and the mass (kDa) is represented by numbers above the peaks. $BrxC^{Eferg}$ displayed a higher ratio of the early eluting peak relative to the dimer peak compared to that observed for $BrxC^{Aci}$ (**Figure 1d**). The $BrxC^{Eferg}$(E268Q) Walker B mutant, expected to allow ATP binding but not hydrolysis, displayed a similar SEC elution profile as $BrxC^{Eferg}$ WT.

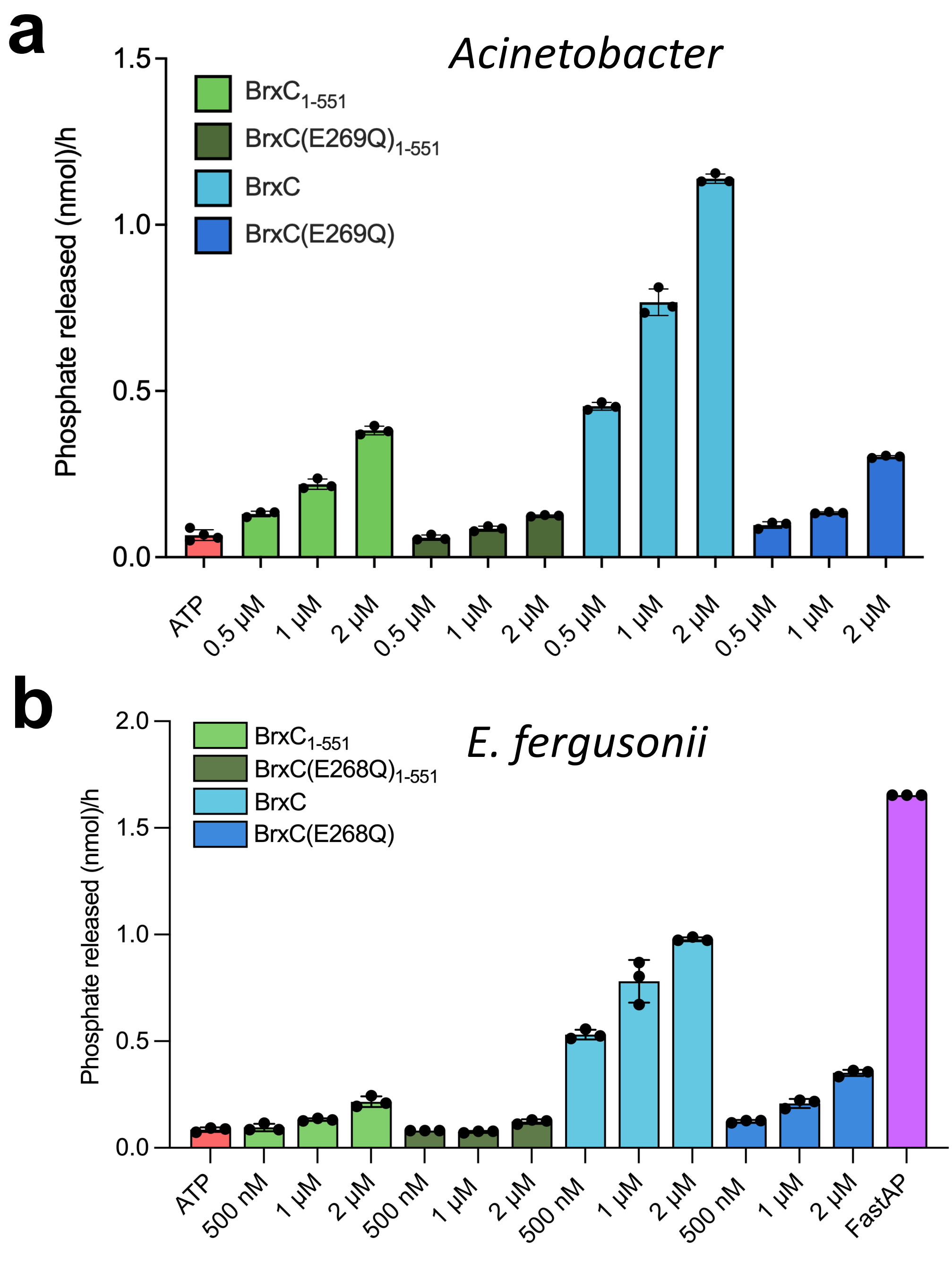


**Supplementary Figure S3. BrxC ATPase activity**.

***Panel a:*** ATPase activity of *Acinetobacter* BrxC and mutants. The phosphate released during a 1 hr reaction containing 500 nM, 1 µM, and 2 µM $BrxC^{Aci}$, $BrxC^{Aci}$(E269Q), $BrxC^{Aci}_{1\text{-}551}$, or $BrxC^{Aci}(E269Q)_{1\text{-}551}$, in the presence 100 µM ATP and reaction buffer containing 10 mM $MgCl_2$ incubated at 37 °C. Control reactions contained 100 µM ATP.

***Panel b:*** ATPase activity of *E. fergusonii* BrxC and mutants. The phosphate released during a 1 hr reaction containing 500 nM, 1 µM, and 2 µM BrxC, $BrxC^{Eferg}$(E268Q), $BrxC^{Eferg}_{1\text{-}551}$, or $BrxC^{Eferg}(E268Q)_{1\text{-}551}$, in the presence 100 µM ATP and reaction buffer containing 10 mM $MgCl_2$ incubated at 37 °C. Control reactions contained 100 µM ATP or 1 U FastAP thermosensitive alkaline phosphatase in reaction buffer. For both panels, reaction buffer was used as a blank. Triplicate data points are shown. Error bars represent standard deviation. The phosphate released was calculated using standards containing free phosphate which were serially diluted to obtain a slope intercept form to allow for calculations from experimental results.

### Supplementary Figure S4

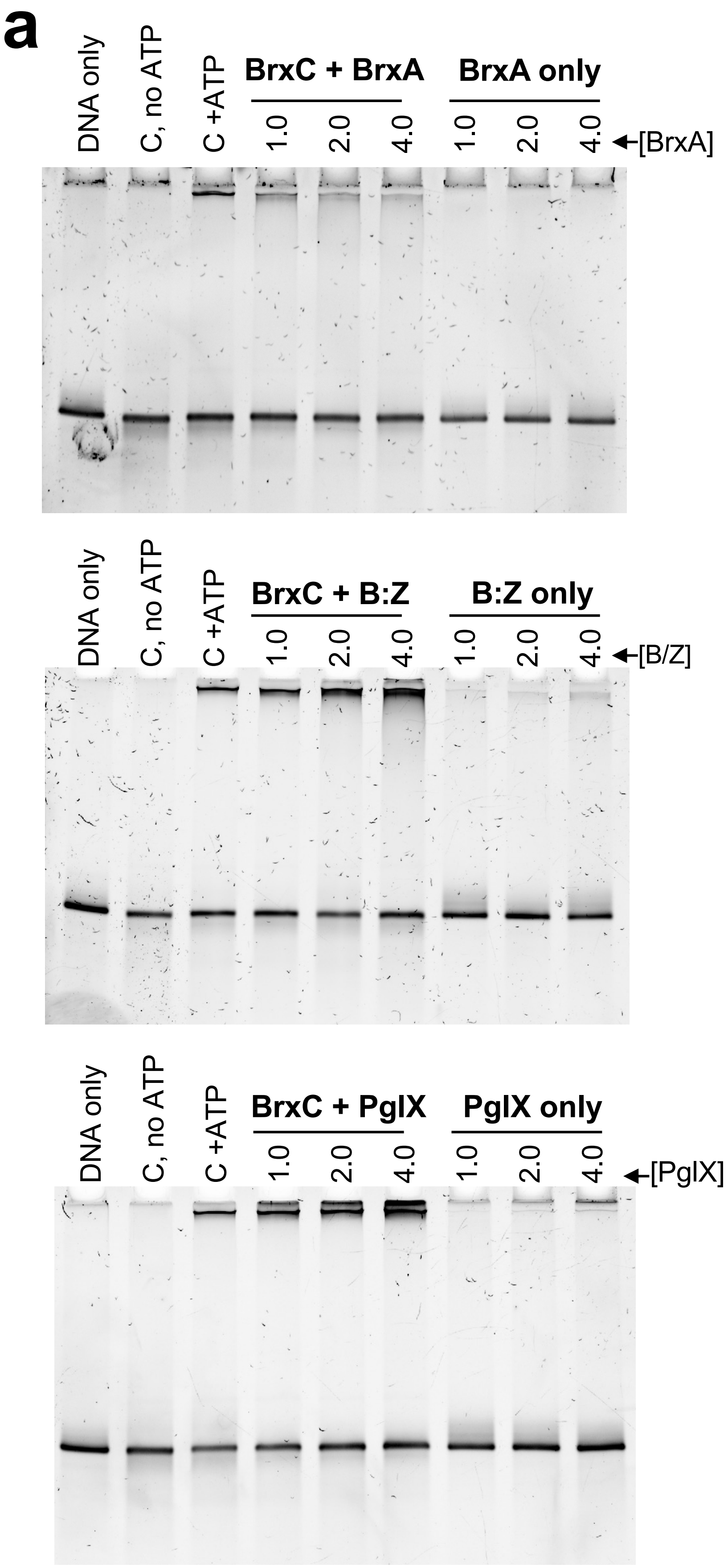


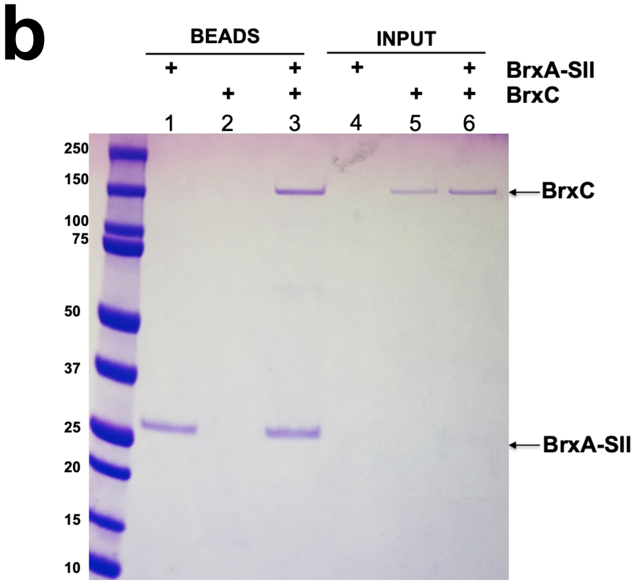


**Supplementary Figure S4. Effect of purified *Acinetobacter* BREX proteins, alone or in combination, on BrxC's interaction with DNA.**

***Panel a:*** BrxC$^{Aci}$-DNA EMSA analyses in the presence of (1) BrxA$^{Aci}$, (2) a tightly associated complex of BrxB$^{Aci}$ and PglZ$^{Aci}$ ("B:Z"), and (3) PglX$^{Aci}$. 2 µM BrxC$^{Aci}$ was incubated with 1, 2 or 4 µM of the indicated proteins for 2 hr at 30 °C prior to separation on native PAGE and visualization of DNA. Incubation of BrxC$^{Aci}$ with B$^{Aci}$:Z$^{Aci}$ or PglX$^{Aci}$ individually does not inhibit DNA binding; in contrast, BrxA$^{Aci}$ reduces BrxC$^{Aci}$ DNA binding.

***Panel b:*** Interaction analysis of BrxA$^{Aci}$ and BrxC$^{Aci}$ via a pull-down experiment with untagged BrxC$^{Aci}$ and BrxA$^{Aci}$ harboring a C-terminal twin-strep affinity tag ("BrxA-SII").

### Supplementary Figure S5

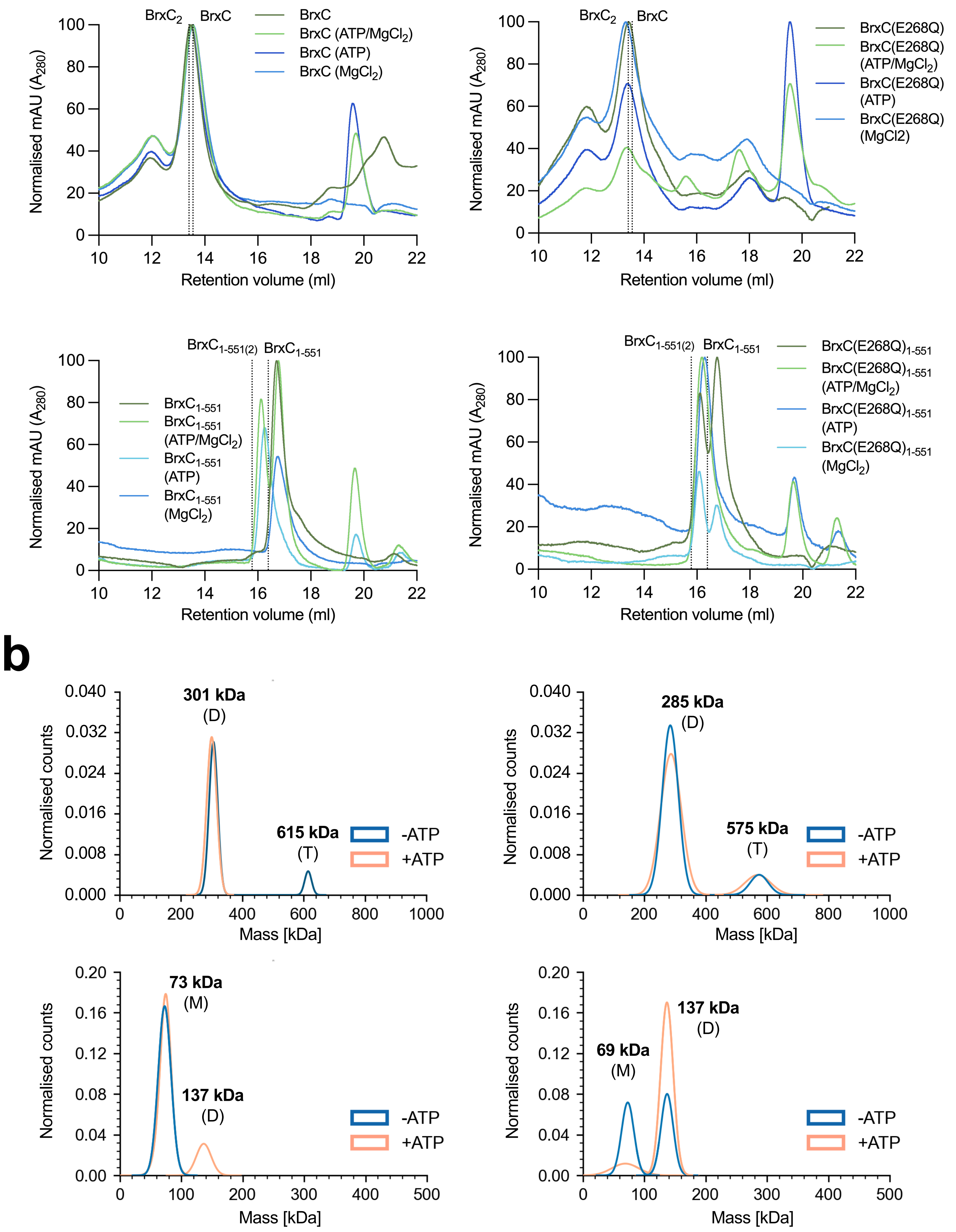

**Supplementary Figure S5. BrxC from *E. fergusonii* undergoes ATP-dependent dimerization.**

***Panel a:*** Analytical size exclusion chromatography using a Superdex 200 increase 10/300 GL column. $BrxC^{Eferg}$, $BrxC^{Eferg}$(E268Q), $BrxC^{Eferg}_{1-551}$, and $BrxC^{Eferg}(E268Q)_{1-551}$, were incubated at 10 μM in the presence and absence of 1 mM ATP and 10 mM $MgCl_2$ for 1 hr at 37 °C or at RT. The expected elution volumes of the monomeric and dimeric proteins are shown by black dotted lines. The traces represent the normalized absorbance at 280 nM. Identifying whether full-length $BrxC^{Eferg}$ and $BrxC^{Eferg}$(E268Q) were monomeric or dimeric was not possible by analytical-SEC based on the expected retention volumes. The solution state was unchanged with the addition of $MgCl_2$ and/or ATP. Additionally, an early eluting peak can be observed on both $BrxC^{Eferg}$ and $BrxC^{Eferg}$(E268Q) analytical-SEC traces with a retention volume of 12 ml, possibly corresponding to a 615 kDa/575 kDa protein or tetrameric $BrxC^{Eferg}$ (556 kDa). Truncated $BrxC^{Eferg}_{1-551}$ exists in solution predominantly as a monomer (63 kDa) with a retention volume of ~17 ml.

***Panel b:*** Mass photometry analysis of $BrxC^{Eferg}$, $BrxC^{Eferg}$(E268Q), $BrxC^{Eferg}_{1-551}$, and $BrxC^{Eferg}(E268Q)_{1-551}$, positioned as per panels in (a). Both dimeric and tetrameric forms were observed for full-length $BrxC^{Eferg}$ WT and $BrxC^{Eferg}$(268Q), supporting the traces seen by analytical SEC. Truncated $BrxC^{Eferg}_{1-551}$ formed dimers in an ATP-dependent manner, and dimer formation was stabilized in the E268Q mutant, indicating ATP hydrolysis promotes dissociation. Samples at 100 nM were incubated with and without 1 μM ATP in the presence of 10 μM $MgCl_2$ for 1 hr at 37 °C. Prior to analysis, samples were diluted directly on the slide by adding 1 μl protein to 9 μl PBS. Data were obtained in the form of mass photometry videos recorded for one minute. Gaussian fits were generated for each data set and plotted on graphs showing the normalised counts against the mass in kDa. The measured mass (kDa) of the monomer (M), dimer (D), and tetramer (T) are shown.

### Supplementary Figure S6

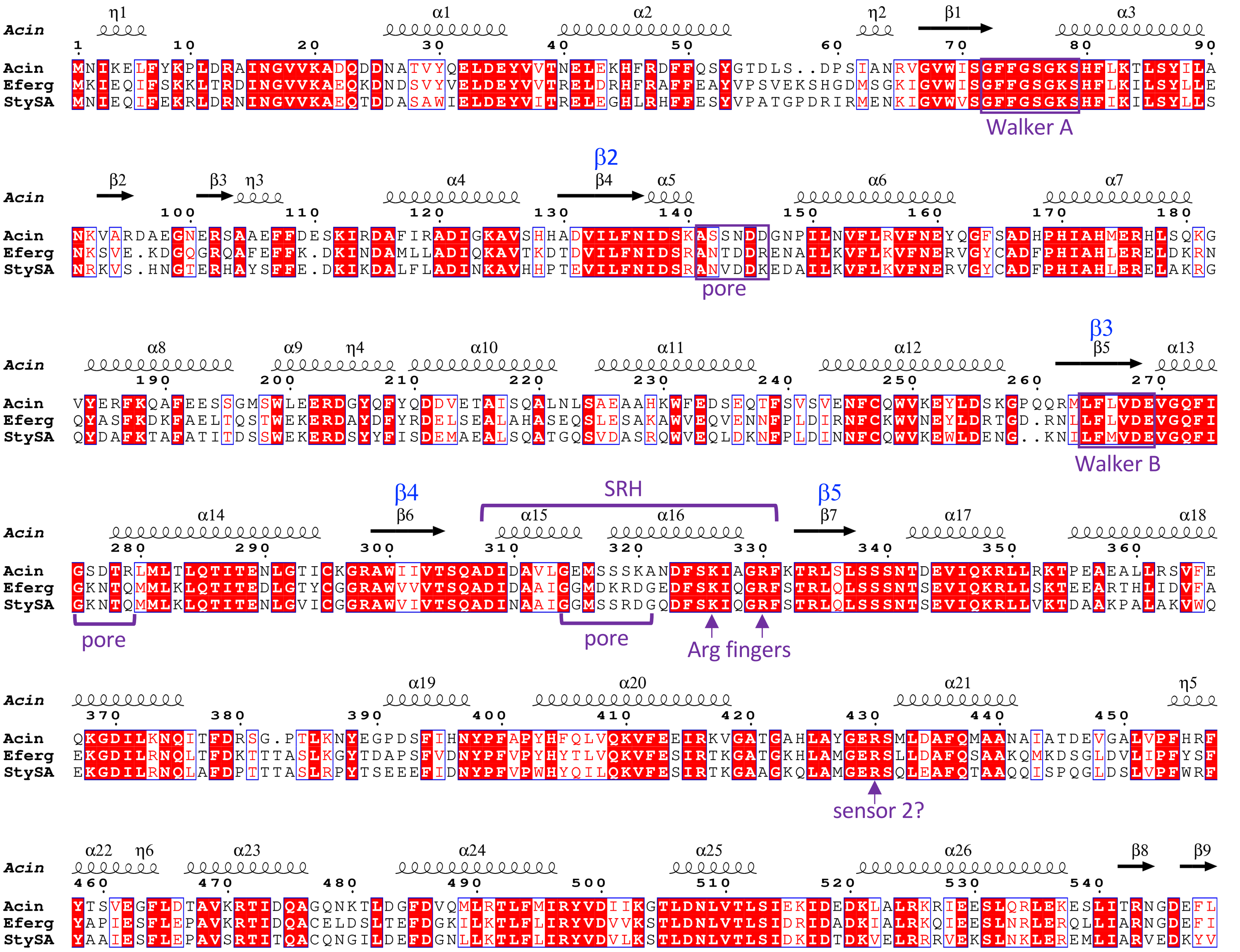


| Motif | Key residues | BrxC[Aci] residue | Typical Mutations | Typical Effects | BrxC mutants | Effect on Dimerization (MP) |
|---|---|---|---|---|---|---|
| Walker A | GxxxxGK[TS] | K79 | K→A | Inhibits nucleotide binding | | |
| Walker B | E in hhhhDE | E269 | E→Q | Impairs ATP hydrolysis; often "**substrate trap**". | E269Q | increase |
| Sensor 1 | Polar residue end of β4 | Q305 | N/T→A | Impairs ATP hydrolysis | | |
| Arg fingers | R or K in the SRH at the end of α4. Often there are 2, sometimes one. | K326, R330 | R→E | Impairs ATP hydrolysis and oligomerization | K326E<br>R330E | decrease<br>decrease |
| Sensor 2 | R near N-term of α7 | R430 | R→A/M | Impairs ATP hydrolysis | | |
| Pore loop | YVG in β2-α2 loop | aa 141-146 | | Impairs substrate binding | | |

**Supplementary Figure S6. Conservation and role of key residues in BrxC AAA+ domain.** ***Top***: Sequence alignment of BrxC AAA+ domains from *Acinetobacter sp. 394*, *Escherichia fergusonii*, and *Salmonella* Typhimurium. The alignments were generated using Clustal Omega, and Espript3.2 was used to indicate secondary structural elements based on the cryo-EM structure of $BrxC^{Aci}$. Conserved AAA+ residues defined in (36) are indicated on the figure; b-strands (labeled in blue) conform to the numbering scheme in reference (36). SRH, second region of homology. ***Bottom***: Motifs, residues and mutations described in this study are based on the BrxC protein sequence from *Acinetobacter sp. 394.*

### Supplementary Figure S7

a

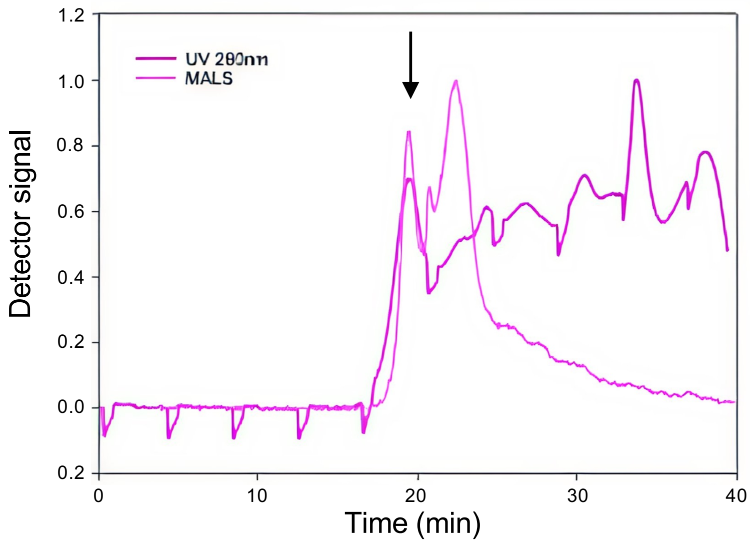


b

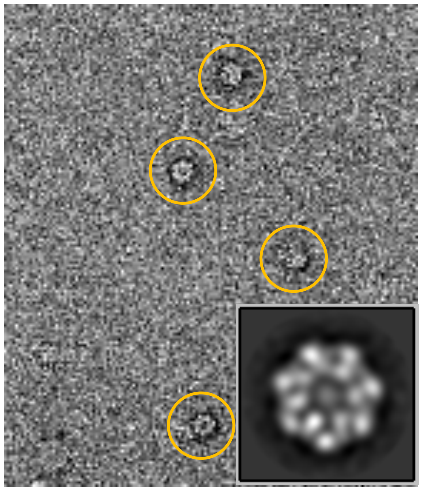

**Supplementary Figure S7. Negative-stain transmission electron micrograph (TEM) of the BrxC complex from *E. fergusonii.***

***Panel a:*** SEC-MALS analysis of the BrxC$^{Eferg}$ complex. BrxC$^{Eferg}$ was separated using a Superdex 200 Increase 10/300 GL column coupled in-line with a multi-angle light scattering (MALS) detector. The chromatogram shows the UV absorbance at 280 nm and the corresponding molar mass distribution across the elution profile, as determined by MALS. The peak selected for TEM analysis is indicated by the black arrow.

***Panel b:*** Representative micrograph from a dataset of 495 negative-stain images, showing BrxC$^{Eferg}$ complexes. Inset, reference-free 2D average obtained after image processing, highlighting the overall shape and symmetry of the complex.

**Supplementary Figure S8**

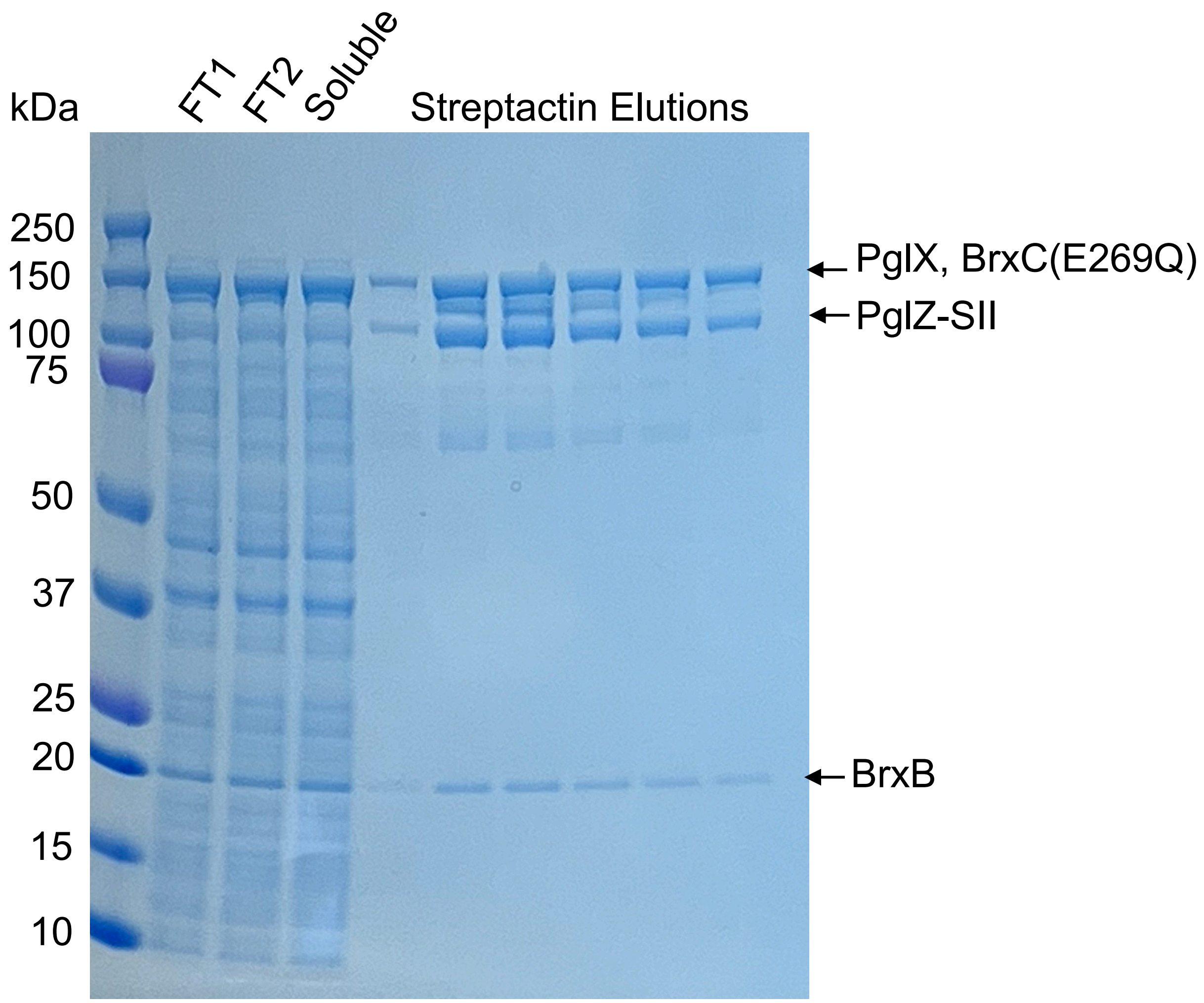


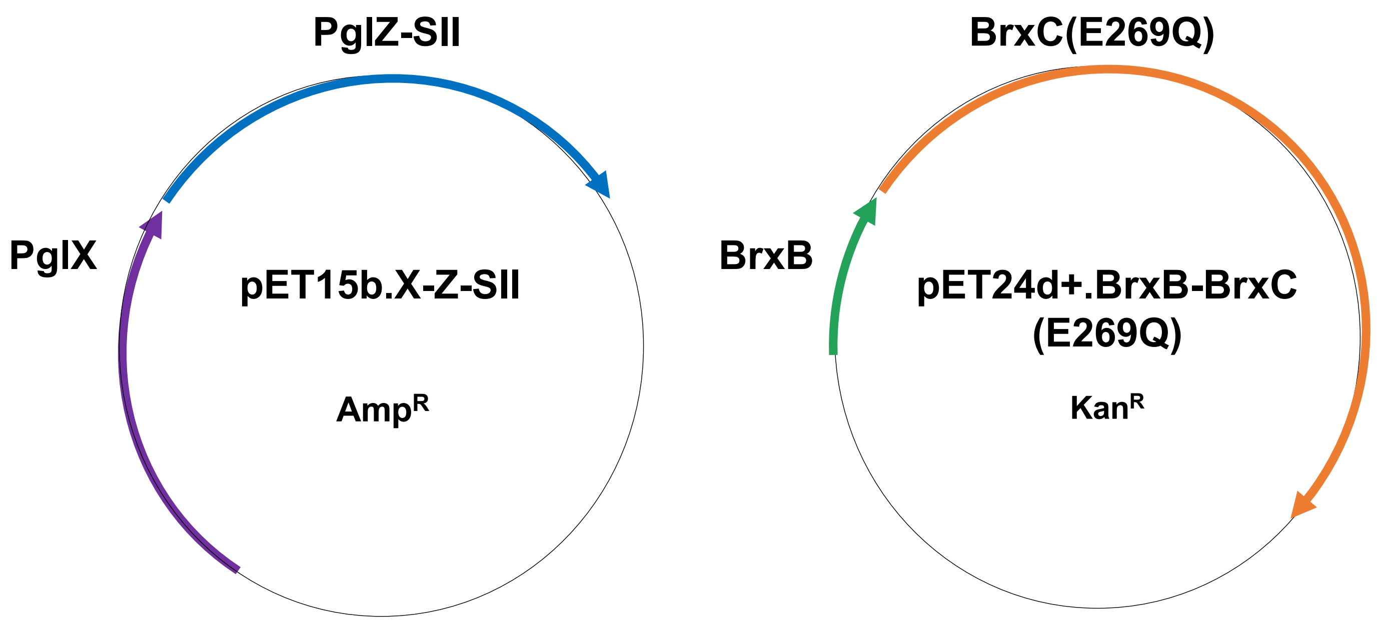

**Supplementary Figure S8. Formation of an *Acinetobacte* BREX complex containing BrxB, PglZ, BrxC and PglX.** We observed the association of *Acinetobacter* $BrxB^{Aci}$, $PglZ^{Aci}$, $BrxC^{Aci}$(E269Q) and $PglX^{Aci}$ (with $PglZ^{Aci}$ containing a twin-strep affinity tag) when all four factors were co-expressed from a pair of plasmids (bottom) in *E. coli* and affinity purified by streptactin resin*,* consistent with observations in other BREX systems (21,22). The identity of $PglX^{Aci}$ and $BrxC^{Aci}$, which overlap on SDS PAGE, were confirmed by mass spectroscopy.

**Supplementary Figure S9**

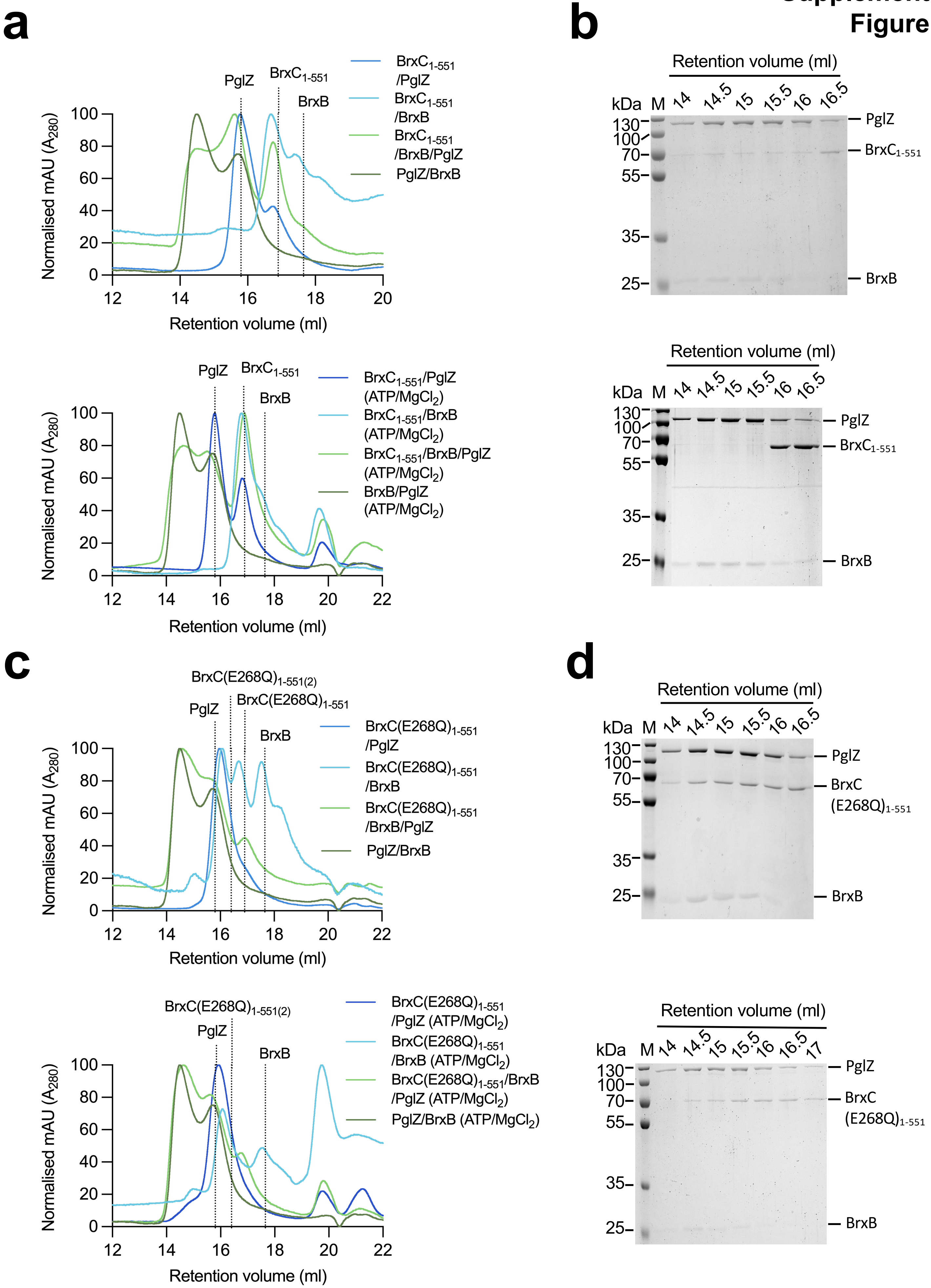

**Supplementary Figure S9. Interactions visualized between *E. fergusonii* BrxB, BrxC and PglZ.**

***Panel a:*** Analytical SEC of *E. fergusonii* BREX proteins using a Superdex 200 increase 10/300 GL column. $BrxC^{Eferg}_{1\text{-}551}$, and $BrxC^{Eferg}(E268Q)_{1\text{-}551}$, were incubated at 10 µM with and without 10 µM $BrxB^{Eferg}$ and 10 µM $PglZ^{Eferg}$, in the presence and absence of 1 mM ATP and 10 mM $MgCl_2$ for 1 hr at 37 °C. The expected elution volume of $PglZ^{Eferg}$, $BrxB^{Eferg}$, and $BrxC^{Eferg}_{1\text{-}551}$, $BrxC^{Eferg}$(E268Q), and $BrxC^{Eferg}$(E268Q) dimer are shown by black dotted lines. The traces represent the normalized absorbance at 280 nM. 0.5 ml fractions were collected.
***Panel b:*** SDS-PAGE analysis (12% polyacrylamide) of fractions collected from analytical SEC. 20 µl of peak fractions were mixed with 10 µl 3x SDS loading dye, boiled for 5 min, and loaded onto the gel. Samples were resolved in 1x tris-glycine running buffer at 180 V for 1 hr 15 min alongside PageRuler™ Plus Prestained protein ladder. Gels were stained with Coomassie Brilliant Blue for 1 hr and destained in MiliQ for 1 hr. Gel images were obtained on a ChemiDoc™ Imaging System. The molecular weights (kDa) of the marker (M) are shown. The expected location of the proteins on the gel are indicated.

**Supplementary Figure S10**

**a**

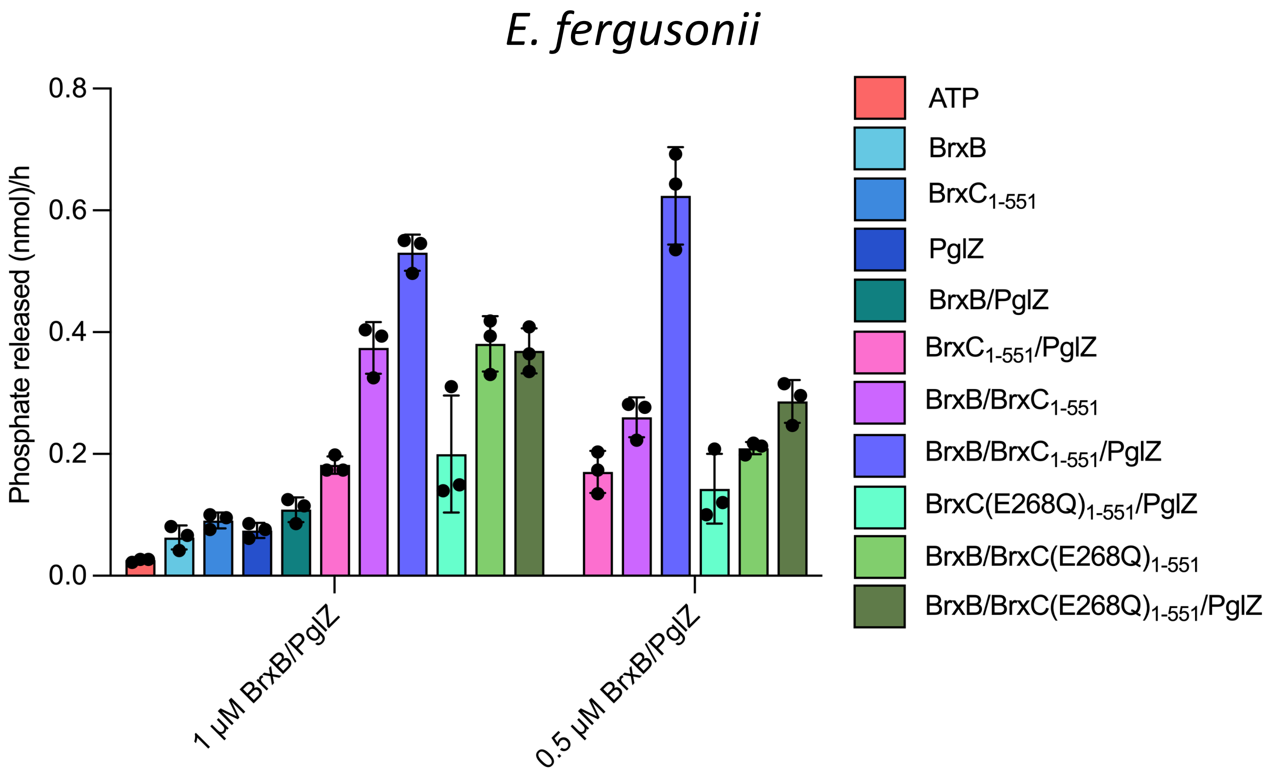


**b**

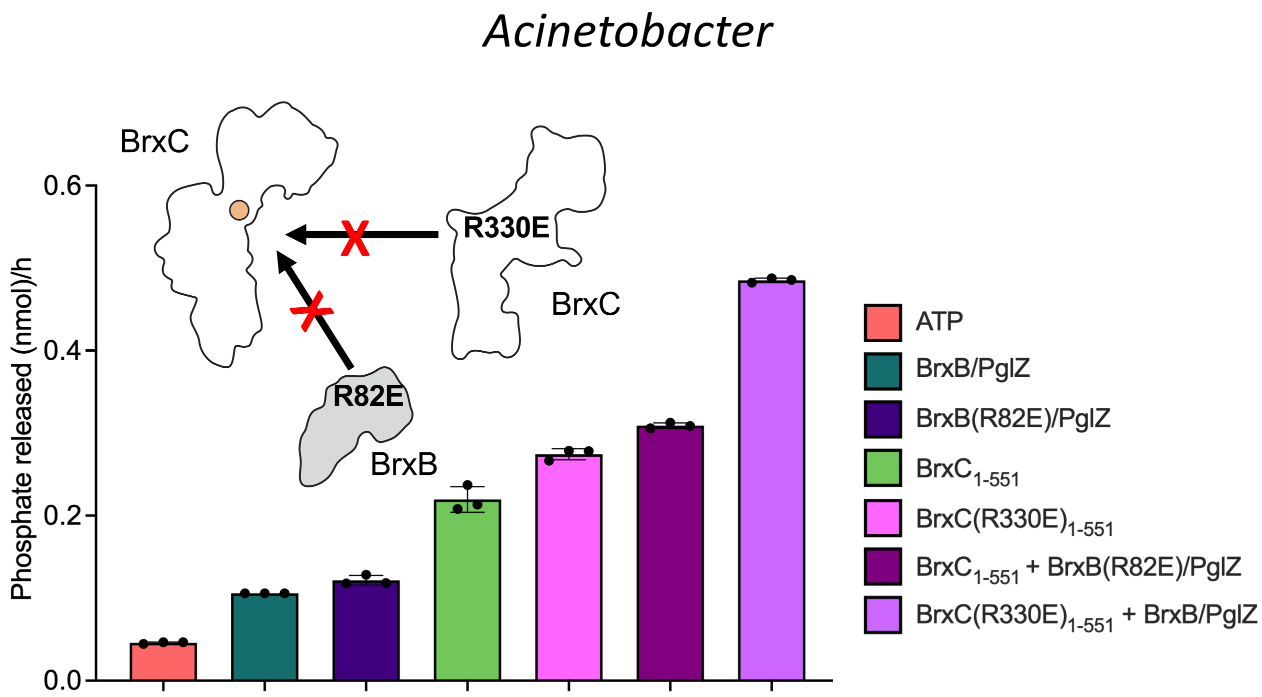

**Supplementary Figure S10. Impacts of BrxB and PglZ on BrxC ATPase activity.**

***Panel a:*** ATPase activity of *E. fergusonii* BrxC mutants and binding partners. The phosphate released during a 1 hr reaction containing 1 µM $BrxC^{Eferg}_{1\text{-}551}$ or $BrxC^{Eferg}(E268Q)_{1\text{-}551}$, with and without 0.5 µM or 1 µM $BrxB^{Eferg}$ and $PglZ^{Eferg}$, in the presence of 10 mM $MgCl_2$ and 100 µM ATP, incubated at 37 °C. Control reactions contained 100 µM ATP or 1 µM protein only in reaction buffer. Reaction buffer was used as a blank. Triplicate data points are shown. Error bars represent standard deviation. The phosphate released was calculated using standards containing free phosphate which were serially diluted to obtain a slope intercept form to allow for calculations from experimental results.

***Panel b:*** ATPase activity of *Acinetobacter* BrxC mutants and binding partners. The phosphate released during a 1 hr reaction containing 1 µM $BrxC^{Aci}(R330E)_{1\text{-}551}$ and $BrxB^{Aci}(R82E)$ with or without 1 µM $BrxB^{Aci}$:$PglZ^{Aci}$ or $BrxC^{Aci}_{1\text{-}551}$ in the presence of 10 mM $MgCl_2$ and 100 µM ATP, incubated at 37 °C and processed and presented as described above. Data presented for $BrxC^{Aci}_{1\text{-}551}$ (green) were taken from **Supplementary Figure S3a** with values calculated internally from that experiment.

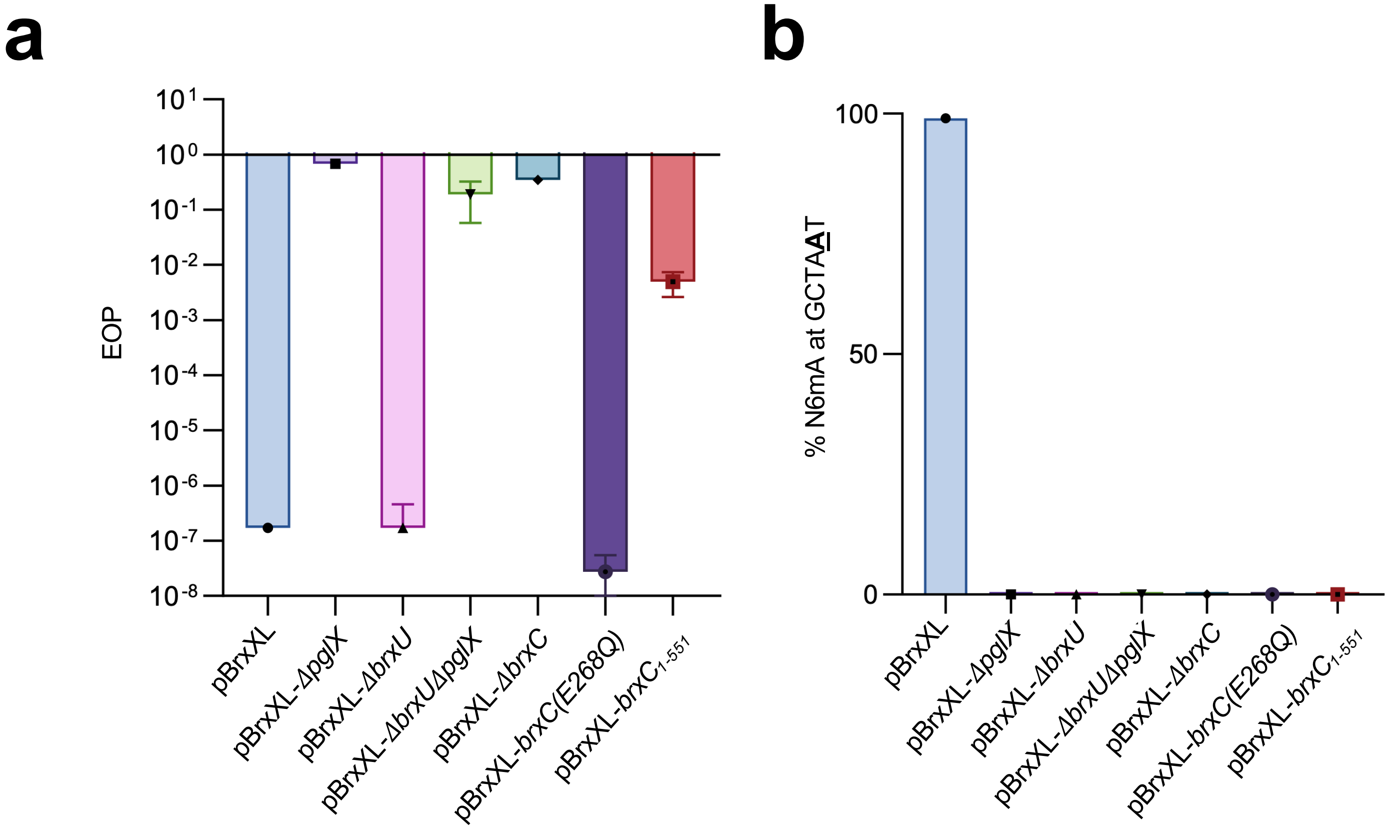


**Supplementary Figure S11. Analysis of *E. fergusonii* BREX phenotypes.**

***Panel a:*** *E. fergusonii* Efficiency of Plating (EOP) data using phage Pau. Mutations of $BrxC^{Eferg}$ were made in the context of the BREX WT plasmid pBrxXL and are shown alongside controls pBrxXL-*ΔpglX* (BREX-), pBrxXL-*ΔbrxU* (BREX+) and pBrxXL-*ΔbrxUΔpglX* (BREX-). The tested mutants of BrxC were Δ*brxC, brxC*(E268Q) and *brxC*$_{1\text{-}551}$. Deletion of *brxC* and truncation 1-551 impairs BREX defence, while $BrxC^{Eferg}$(E268Q) does not cause an impact.

***Panel b:*** PacBio methylation data of the samples tested by EOP. Sequencing percentage corresponds to the amount of methylated GCTA**A**T N6mA BREX motifs. There was no methylation of the $BREX^{Eferg}$ motif across all tested BrxC mutants.
