## Supplemental Tables for "Competing forms of protein-protein association and DNA binding exhibited by BrxC from the BREX phage restriction system"

### SUPPLEMTARY TABLE S1. PLASMIDS USED IN THIS STUDY

| Acin BREX Constructs | Name | Vector | Construct | Purification | Tag/Terminus | Protease | Induction: Auto/IPTG | Antibiotic |
|---|---|---|---|---|---|---|---|---|
| | SII-TEV-BrxC FL WT | pET15b | BrxC FL WT | Streptactin | N-terminal | TEV | Auto | Amp |
| | SII-TEV-BrxC(E269Q) FL | pET15b | BrxC(E269Q) | Streptactin | N-terminal | TEV | Auto | Amp |
| | SII-TEV BrxC(D22C) FL | pET15b | BrxC(D22C) | Streptactin | N-terminal | TEV | Auto | Amp |
| | BrxC(1-1146)-Thr-SII | pET15b | BrxC(1-1147) WT | Streptactin | C-Terminal | Thrombin (not used) | IPTG | Amp |
| | BrxC(1-1146)E269Q-Thr-SII | pET15b | BrxC(1-1147)E269Q | Streptactin | C-Terminal | Thrombin (not used) | IPTG | Amp |
| | BrxC(1-551)-TEV-SII WT | pET15b | BrxC(1-551) WT | Streptactin | C-Terminal | TEV | IPTG | Amp |
| | BrxC(1-551)E269Q-TEV-SII | pET15b | BrxC(1-551)E269Q | Streptactin | C-Terminal | TEV | IPTG | Amp |
| | BrxC(1-551)R330E-TEV-SII | pET15b | BrxC(1-551)R330E | Streptactin | C-Terminal | TEV | IPTG | Amp |
| | PglZ-Thr-SII | pET15b | PglZ WT | Streptactin | C-Terminal | Thrombin | IPTG | Amp |
| | PglZ-Thr-SII | pET24d | PglZ WT | Streptactin | C-Terminal | Thrombin | IPTG | Kan |
| | PglZ(1-98)-Thr-SII | pET15b | PglZ(1-98) | Streptactin | C-Terminal | Thrombin | IPTG | Amp |
| | BrxB | pET24d | BrxB WT | Copurified | N/A | N/A | IPTG | Kan |
| | BrxB(R82E) | pET24d | BrxB(R82E) | Copurified | N/A | N/A | IPTG | Kan |
| | BrxB(S152C) | pET24d | BrxB(S152C) | Copurified | N/A | N/A | IPTG | Kan |
| | SII-TEV-PglX | pET15b | PglX WT | Streptactin | N-terminal | TEV | Auto | Amp |
| | BrxA-Thr-SII | pET15b | BrxA WT | Streptactin | C-Terminal | Thrombin | IPTG | Amp |
| | BrxC | pET24d | BrxC No Tag | Copurified | N/A | N/A | IPTG | Kan |
| | BrxB + BrxC | pET24d | BrxB + BrxC WT | Copurified | N/A | N/A | IPTG | Kan |
| | BrxB + BrxC(E269Q) | pET24d | BrxB + BrxC(E269Q) | Copurified | N/A | N/A | IPTG | Kan |
| | BrxB + BrxC(1-551) | pET24d | BrxB + BrxC(1-551) | Copurified | N/A | N/A | IPTG | Kan |
| | BrxA + BrxB + BrxC(E269Q) | pET24d | BrxA + BrxB + BrxC WT | Copurified | N/A | N/A | IPTG | Kan |
| | PglX + PglZ-SII | pET15b | PglX + PglZ-SII | Sreptactin | C-Terminal | Not Used | IPTG | Amp |
| | SII-TEV-PglZ + PglZ | pET15b | PglX + PglZ | Streptactin | N-Terminal | Not Used | IPTG | Amp |
| | H6-BrxC(1-553)-BrxB Fusion | pET15b | rxC(1-553)-BrxB Fusio | Co-purified | His6 - N-Termina | N/A | IPTG | Amp |
| | pACYC BREX WT | pACYC | WT BREX | N/A | N/A | N/A | N/A | Chlor |
| | pACYC BREX ΔBrxC | pACYC | BREX del.BrxC | N/A | N/A | N/A | N/A | Chlor |
| | pACYC BREX ΔBrxB | pACYC | BREX del.BrxB | N/A | N/A | N/A | N/A | Chlor |
| | pACYC BREX BrxC(E269Q) | pACYC | BREX + BrxC(E269Q) | N/A | N/A | N/A | N/A | Chlor |
| | pACYC BREX BrxC(R330E) | pACYC | BREX + BrxC(R330E) | N/A | N/A | N/A | N/A | Chlor |
| | pACYC BREX BrxC(1-1146) WT | pACYC | BREX + BrxC(1-1146) | N/A | N/A | N/A | N/A | Chlor |
| | pACYC BREX BrxC(1-1551) WT | pACYC | BREX + BrxC(1-551) | N/A | N/A | N/A | N/A | Chlor |
| | pACYC BREX BrxB(R82E) | pACYC | BREX + BrxB(R82E) | N/A | N/A | N/A | N/A | Chlor |
| | pACYC BREX BrxB(R185E) | pACYC | BREX + BrxB(R185E) | N/A | N/A | N/A | N/A | Chlor |
| **E. ferg BREX Constructs** | pTRB791 | pSAT1-LIC | BrxC | IMAC | -SUMO - N-Term | hSENP2 | Auto | Amp/Chlor |
| | pTRB792 | pSAT1-LIC | BrxC(E268Q) | IMAC | -SUMO - N-Term | hSENP2 | Auto | Amp/Chlor |
| | pTRB789 | pSAT1-LIC | BrxC(1-551) | IMAC/AEC | -SUMO - N-Term | hSENP2 | IPTG | Amp/Tet |
| | pTRB790 | pSAT1-LIC | BrxC(1-551)E268Q | IMAC/AEC | -SUMO - N-Term | hSENP2 | IPTG | Amp/Tet |
| | pTRB449 | pSAT1-LIC | PglZ | IMAC/AEC | -SUMO - N-Term | hSENP2 | IPTG | Amp |
| | pTRB444 | pSAT1-LIC | BrxB | IMAC/AEC | -SUMO - N-Term | hSENP2 | IPTG | Amp |
| | pBrxXL | pBrxXL | WT BREX | N/A | N/A | N/A | N/A | Chlor |
| | pBrxXL-ΔPglX | pBrxXL | BREX del.PglX | N/A | N/A | N/A | N/A | Chlor |
| | pBrxXL-ΔBrxU | pBrxXL | BREX del.BrxU | N/A | N/A | N/A | N/A | Chlor |
| | pBrxXL-ΔBrxUΔPglX | pBrxXL | BREX del.BrxU/PglX | N/A | N/A | N/A | N/A | Chlor |
| | pBrxXL-ΔBrxC | pBrxXL | BREX del.BrxC | N/A | N/A | N/A | N/A | Chlor |
| | pBrxXL-BrxC(E268Q) | pBrxXL | BREX + BrxC(E268Q) | N/A | N/A | N/A | N/A | Chlor |
| | pBrxXL-BrxC(1-551) | pBrxXL | BREX + BrxC(1-551) | N/A | N/A | N/A | N/A | Chlor |

| | | | |
|---|---|---|---|
| **dsDNA Substrates** | 100 bp PglX target centered | N/A | gcaggatcgaatcatggctggcgtcatgtaactacaggaatggtatg**GTAGAT**gatcgcatagattgatgcatacaatccgtttacggattcaggtaccg |

**SUPPLEMENTARY TABLE S2. CryoEM Data**

| | BrxC-E269Q | BrxCBZ dimer | BrxCBZ |
|---|---|---|---|
| PDB/EMD-ID | 9ZDX/EMD-74076 | 9ZN5/EMB-74435 | EMD-74400 |
| *Data collection* | | | |
| EM equipment | Glacios | Glacios | Glacios |
| Voltage (kV) | 200 | 200 | 200 |
| Detector | Gatan K3 | Gatan K3 | Gatan K3 |
| Pixel size (Å/Pixel) | 0.561 | 0.561 | 0.561 |
| Electron dose (e-/Å^2) | 50 | 50 | 50 |
| Defocus range (um) | | | |
| Micrographs collected | 4131 | 4012 | 3965 |
| Micrographs used | 3686 | 2509 | 3602 |
| *Reconstruction* | | | |
| Software | cryoSPARC | cryoSPARC | cryoSPARC |
| # particles used | 477,102 | 195,504 | 101,355 |
| Symmetry | None | None | None |
| Resolution (Å) | 2.95 | 7.02 | 4.46 |
| Map sharpening B-factor (Å) | 116.7 | 677.9 | 254.9 |
| *Refinement* | | | |
| Software | phenix | N/A | N/A |
| Box | | | |
| Lengths (Å) | 79.7, 96.5, 131.3 | N/A | N/A |
| Angles (°) | 90.0, 90.0, 90.0 | N/A | N/A |
| *Model composition* | | | |
| Protein residues | 1094 | N/A | N/A |
| Nucleotide | 0 | N/A | N/A |
| Water | 4 | N/A | N/A |
| Ligands | 4 | N/A | N/A |
| MolProbity score | 1.53 | N/A | N/A |
| RMS Bonds (Å) | 0.004 | N/A | N/A |
| RMS Angle (°) | 0.539 | N/A | N/A |
| Ramachandran favored (%) | 96.51 | N/A | N/A |
| Ramachandran allowed (%) | 3.49 | N/A | N/A |
| Ramachandran outliers (%) | 0 | N/A | N/A |

**Supplementary Table S3. BrxC$_{1\text{-}551}$:BrxB:PglZ$_{1\text{-}98}$ crystallographic data and refinement**

**PDB code**: 9ZLL

| **Data Collection and Processing** | | |
|---|---|---|
| Unit cell Dimensions | 149.25 114.99 194.05 90.00 108.03 90.00 | |
| Space Group | I2 | |
| Average Mosaicity | 0.17 ° | |
| | Overall | Outer Shell |
| Low resolution limit | 49.68 | 2.79 |
| High resolution limit | 2.74 | 2.74 |
| Rmerge | 0.085 | 0.711 |
| Rpim | 0.038 | 0.318 |
| Total number of observations | 555957 | 29943 |
| Total number unique | 81941 | 4446 |
| Mean((I)/$\sigma$(I)) | 14.3 | 2.2 |
| Mn(I) half-set correlation CC(1/2) | 0.998 | 0.782 |
| Completeness (%) | 99.9 | 99.7 |
| Multiplicity | 6.8 | 6.7 |

| **Model refinement** | |
|---|---|
| $R_{work}$ | 0.2195 |
| $R_{free}$ | 0.2674 |
| Mean Biso | 69.1 Å$^2$ |
| $RMSD_{bond}$ | 0.005 Å |
| $RMSD_{angles}$ | 0.98° |
| $RMSD_{dihedral}$ | 15.337° |
| Ramachandran distribution (Favored, allowed, disallowed) | 92.85%, 5.54%, 1.61% |
